# Rapid cortical reorganization tracks goal-directed sensorimotor learning in real time

**DOI:** 10.64898/2026.05.11.724293

**Authors:** Anthony Renard, Georgios Foustoukos, Maya Iuga, Pol Bech, Axel Bisi, Robin F. Dard, Sylvain Crochet, Carl C.H. Petersen

## Abstract

Animals can learn to execute a specific action in response to a sensory cue predicting reward availability. How neuronal circuits in the brain change to enable goal-directed sensorimotor transformations remains poorly understood. Here, we developed a barrel cortex-dependent whisker-based detection task in which thirsty mice learned to associate a tactile whisker stimulus with licking for a water reward within a single behavioral session and continued to improve performance over subsequent days. Longitudinal two-photon calcium imaging of layer 2/3 neurons in the whisker-related primary somatosensory barrel cortex (wS1) revealed that the neuronal representation of the whisker deflection reorganized with reward learning. Neurons in wS1 projecting to the whisker-related secondary somatosensory cortex (wS2), but not intermingled neurons projecting to the whisker-related primary motor cortex (wM1), on average increased responsiveness to whisker stimulation upon reward learning. Examining the first day of learning, population decoding tracked learning trial-by-trial with neuronal trajectories mirroring behavior. Interestingly, neurons that gained stimulus responsiveness across training preferentially took part in spontaneous reactivation events during learning. Our results reveal that reward-based learning is accompanied by a reorganization of sensory-evoked responses and spontaneous activity patterns in mouse barrel cortex. In future experiments it will be important to examine causal roles for these learning-related changes in neuronal dynamics.

## Introduction

Adaptive behavior requires the brain to rapidly associate sensory stimuli with appropriate motor responses. While learning-related changes across sensory modalities have been extensively investigated, studies frequently rely on training paradigms extending over days or weeks (e.g. (Kato et al., 2015; Poort et al., 2015)), leaving the dynamics of the formation of the initial association unresolved. It remains unclear whether sensory representations in neocortex reorganize on the timescale at which behavioral changes first emerge or whether neocortical plasticity reflects a slower process requiring days of offline consolidation (Drieu et al., 2025; McClelland et al., 1995). To help address this question, here, we developed a whisker-based detection task in which mice form a novel sensorimotor association within a single behavioral session.

Mice use their array of facial whiskers to obtain tactile information about the presence, location, shape and texture of objects in their immediate vicinity (Bosman et al., 2011; Diamond et al., 2008; Petersen, 2019; Staiger and Petersen, 2021). Each whisker is individually represented by an anatomical unit termed a ‘barrel’ in the primary whisker-related somatosensory barrel cortex (wS1) (Woolsey and Van der Loos, 1970). After an early critical period during the first postnatal week in mice, the anatomical barrel map is fixed, providing an important reference upon which functional plasticity of neuronal representations can be studied. Pioneering studies of the effects of sensory deprivation through whisker trimming revealed important changes in sensory responses and specific alterations in synaptic circuits in wS1 driven by experience-dependent plasticity (Allen et al., 2003; Chittajallu and Isaac, 2010; Gainey et al., 2018; Hardingham et al., 2003; Holtmaat et al., 2006; House et al., 2011; Jacob et al., 2012; Shepherd et al., 2003; Trachtenberg et al., 2002; Wen et al., 2013). More recently, sensory systems neuroscientists have increasingly turned to investigating sensory-guided decision-making tasks requiring the learning of goal-directed sensorimotor transformations typically motivated by reward or punishment (Carandini and Churchland, 2013). Complex tasks, for example whisker-based texture discrimination, aperture discrimination and reversal learning, can take weeks for mice to learn, and the accompanying changes in neuronal representations have been tracked across days through longitudinal imaging (Audette et al., 2019; Banerjee et al., 2020; Buetfering et al., 2022; Chen et al., 2015; Chéreau et al., 2020; Han and Helmchen, 2025; Heimburg et al., 2026; Peron et al., 2015a; Rabinovich et al., 2022). However, the trial-by-trial neural dynamics in relationship to changes in behavioral performance are more difficult to establish across days because of the unknown impact of the intervening period between behavioral sessions lacking neurophysiological measurements. Even one of the most elementary tasks in which thirsty mice learn to lick a water reward spout in response to the detection of a single whisker deflection typically requires many days of training before reaching expert performance. Comparing naïve and expert mice revealed prominent changes in evoked sensory responses across various brain regions including hippocampus (Le Merre et al., 2018), medial prefrontal cortex (Le Merre et al., 2018), dorsolateral striatum (Sippy et al., 2021, 2015), and wS1 (Sachidhanandam et al., 2013; Yamashita and Petersen, 2016). However, comparison across cohorts of naïve and expert mice precludes examining plasticity in individual identified neurons, and can therefore only identify changes in classes of neurons across learning, such as: i) an increased fast whisker-evoked response in direct pathway striatal projection neurons, but not indirect pathway striatal projection neurons (Sippy et al., 2021, 2015); and ii) an increased whisker-evoked sensory response and licking-related activity in neurons in wS1 projecting to secondary whisker somatosensory cortex (wS2), but not in the intermingled wS1 neurons projecting to primary whisker motor cortex (wM1) (Yamashita and Petersen, 2016).

To help better understand the relationship between neuronal dynamics and learning, here we developed a whisker-based detection task in which thirsty mice learn to lick a water reward spout in response to a single whisker deflection within a single behavioral session. Combining longitudinal two-photon calcium imaging of layer 2/3 neurons with trial-by-trial population decoding, we show that reward learning is accompanied by rapid reorganization of population representations tracking behavior, and that neurons recruited by learning are preferentially engaged in spontaneous reactivations.

## Results

### Mice rapidly learn a new whisker–reward association within a single session

To study rapid sensorimotor learning, head-fixed, water-restricted mice were trained on a whisker detection task designed to capture the formation of a new sensorimotor association in the course of a single behavioral session (Figure 1A). Mice first went through an initial auditory pre-training phase during which they were rewarded with a drop of water for licking a spout in response to a brief auditory tone. Auditory pre-training continued for 5-7 days until stable expert performance was reached for at least two days (Days −2 and −1). On Day 0, a brief deflection of the C2 whisker, until then delivered only passively after auditory sessions, was introduced in the task, and whisker training continued for two additional days (Days +1 and +2). To establish that behavioral and neuronal changes were driven by reward, mice were randomly assigned to one of two groups: rewarded mice (R+, n = 19 mice during functional imaging, described later) for which licking within 1 s of C2 whisker stimulation was rewarded, and non-rewarded mice (R-, n = 16 mice during functional imaging, described later) for which licking in response to whisker stimulation was not rewarded. R-mice thus experienced the same stimuli and task context as R+ mice but learned to withhold licking to the whisker stimulus. Auditory trials were interleaved throughout training and remained rewarded in both groups, providing an internal control for task engagement. Catch trials without sensory stimulation were pseudo-randomly interleaved to assess spontaneous lick rates. Licking in catch trials was never rewarded.

**Figure 1.**
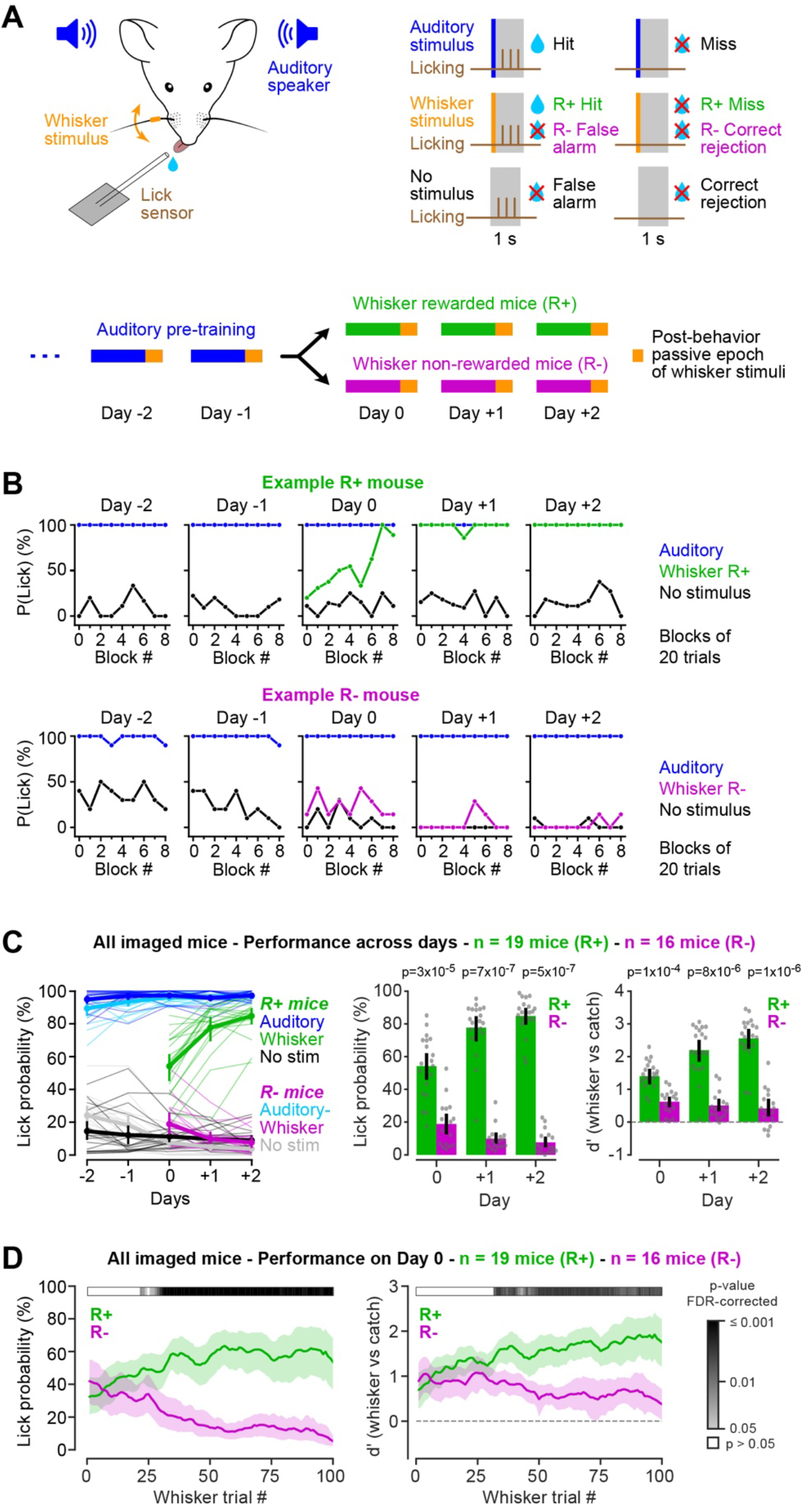
Mice learn to lick for reward in response to a whisker stimulus within a single session. (**A**) Water-restricted mice were first trained to lick a water reward spout in response to a brief auditory pure tone (Days −2 and −1 are the last two days of auditory pre-training). On Day 0, the whisker stimulus (3 ms deflection of the C2 whisker) was introduced as an interleaved trial type, and whisker training continued for two more days (Days +1 and +2). Whisker-rewarded mice (R+) received a water reward if they licked in response to the whisker stimulus (whisker hit), while whisker non-rewarded mice (R-) did not. Auditory hits were rewarded in both groups, and catch trials (no stimulus) were never rewarded. On every day, a block of 50 passive whisker stimuli was delivered after the mouse had disengaged from the task (after three consecutive missed auditory trials). (**B**) Lick probability in blocks of 20 trials across the five days, for one example R+ mouse (top) and one example R-mouse (bottom), on auditory (blue), whisker (green, R+; magenta, R-) and catch trials (black). (**C**) Performance of the imaged mice across days (n = 19 R+ and 16 R-mice). Left: lick probability on auditory (dark blue, R+; light blue, R-), whisker (green, R+; magenta, R-) and catch trials (black, R+; grey, R-). Thin lines, mice; thick lines, mean ± 95% CI. Middle: whisker lick probability on Days 0 to +2. Right: d′ (whisker vs catch trials) on the same days. Bars, mean ± 95% CI; dots, mice; p-values, Mann-Whitney U test, R+ vs R-on each day computed across mice. (**D**) Lick probability (left) and d′ (right) across the first 100 whisker trials of Day 0. For each mouse, lick probability on whisker and catch trials was estimated trial-by-trial with a Bayesian state space model fitted to the binary lick outcomes, and d′ was computed from these two curves (see Methods). Lines, mean ± 95% CI across mice. The grey scale bar above each plot gives the p-value of a Mann-Whitney U test, R+ vs R-, at each whisker trial, corrected for multiple comparisons across trials (Benjamini-Hochberg). White means p > 0.05. The two groups first differed in lick probability at whisker trial 22; they differed in d′ from trial 32 onwards.

Whisker-rewarded mice, but not whisker-unrewarded mice, learned to lick in response to whisker stimulation within a single session (Figures 1B and 1C). The overall probability of licking in response to whisker stimulation on Day 0 was significantly higher in R+ than R-mice (Mann-Whitney U test, n = 19 R+ mice vs 16 R-mice, p = 3x10^-5^; d′, whisker vs catch trials, p = 1x10^-4^) and continued to diverge over the following two days (Figure 1C). Auditory performance remained stable across days in both groups, ruling out general task disengagement as an explanation for the divergent whisker learning trajectories.

On Day 0, the first whisker hit occurred on average within 5 whisker trials (Figure 1–figure supplement 1A). In the first whisker trials of Day 0, both groups licked to the whisker stimulus at similar rates (probability of licking in the first 10 whisker trials: 34% for 19 R+ mice; 38% for 16 R-mice; Mann-Whitney U test, p = 0.81), and more than in catch trials (Wilcoxon signed-rank test: R+, p = 7x10^-4^, n = 19 mice; R-, p = 0.002, n = 16 mice) (Figure 1–figure supplement 1B). This suggests an initial generalization of the auditory lick response to the new stimulus, after which licking was reinforced in R+ mice and suppressed in R-mice. Within Day 0, whisker hit rate progressively increased in R+ mice, while R-mice rapidly learned to suppress licking to the non-rewarded whisker stimulus, with the two groups significantly diverging after 22 whisker trials, 14 ± 4 min into the session (Mann-Whitney U test with Benjamini-Hochberg correction) (Figure 1D and Figure 1–figure supplement 1D-F). Measured as d′ of whisker vs catch, the divergence was significant from whisker trial 32 (Figure 1D). Together, these results demonstrate that mice can rapidly acquire a new stimulus-reward association driven by reward within a single behavioral session, providing a tractable framework for investigating the neural mechanisms underlying fast goal-directed learning.

### Barrel cortex activity is required for whisker-guided licking during learning

Having established that mice rapidly learn the whisker association within a single session, we next asked whether neuronal activity in wS1 might play a role in task acquisition. We therefore carried out inactivation experiments in two different cohorts of whisker-rewarded mice, which were not functionally-imaged.

We first carried out pharmacological inactivation experiments. Muscimol, a GABA-A receptor agonist, was injected into the C2 barrel column of the contralateral wS1 or, as a control, into the forepaw primary somatosensory cortex (fpS1, located ∼1.5 mm anterior to the C2 whisker representation in wS1) (Figure 2A). Inactivation was performed over the first three days of whisker-rewarded learning (Days 0, +1, and +2), followed by three recovery days without injection (Days +3, +4, and +5). During the days of muscimol inactivation in wS1, mice failed to increase their probability of licking in response to whisker stimulation, but mice acquired the task in the subsequent recovery days without muscimol (n = 13 mice) (Figure 2B). In contrast, fpS1 inactivation did not prevent learning (n = 8 mice) (Figure 2B). Whisker-evoked licking probabilities on Day 0 (as well as later days) were significantly lower in mice with wS1 inactivation compared to mice with fpS1 inactivation (Mann-Whitney U test, n = 13 wS1 inactivated mice vs n = 8 fpS1 inactivated mice: Day 0, p = 0.013; Day +1, p = 3x10^-3^; and Day +2, p = 7x10^-4^) (Figure 2C). Similarly, quantifying whisker vs catch lick probability through d’, inactivation of wS1 was found to reduce performance on Day 0 and later days compared to fpS1 inactivated mice (Mann-Whitney U test, n = 13 wS1 inactivated mice vs n = 8 fpS1 inactivated mice: Day 0, p = 3x10^-3^; Day +1, p = 3x10^-3,^ and Day +2, p = 4x10^-5^) (Figure 2C). Auditory performance was not affected by wS1 or fpS1 inactivation (Figure 2–figure supplement 1).

**Figure 2.**
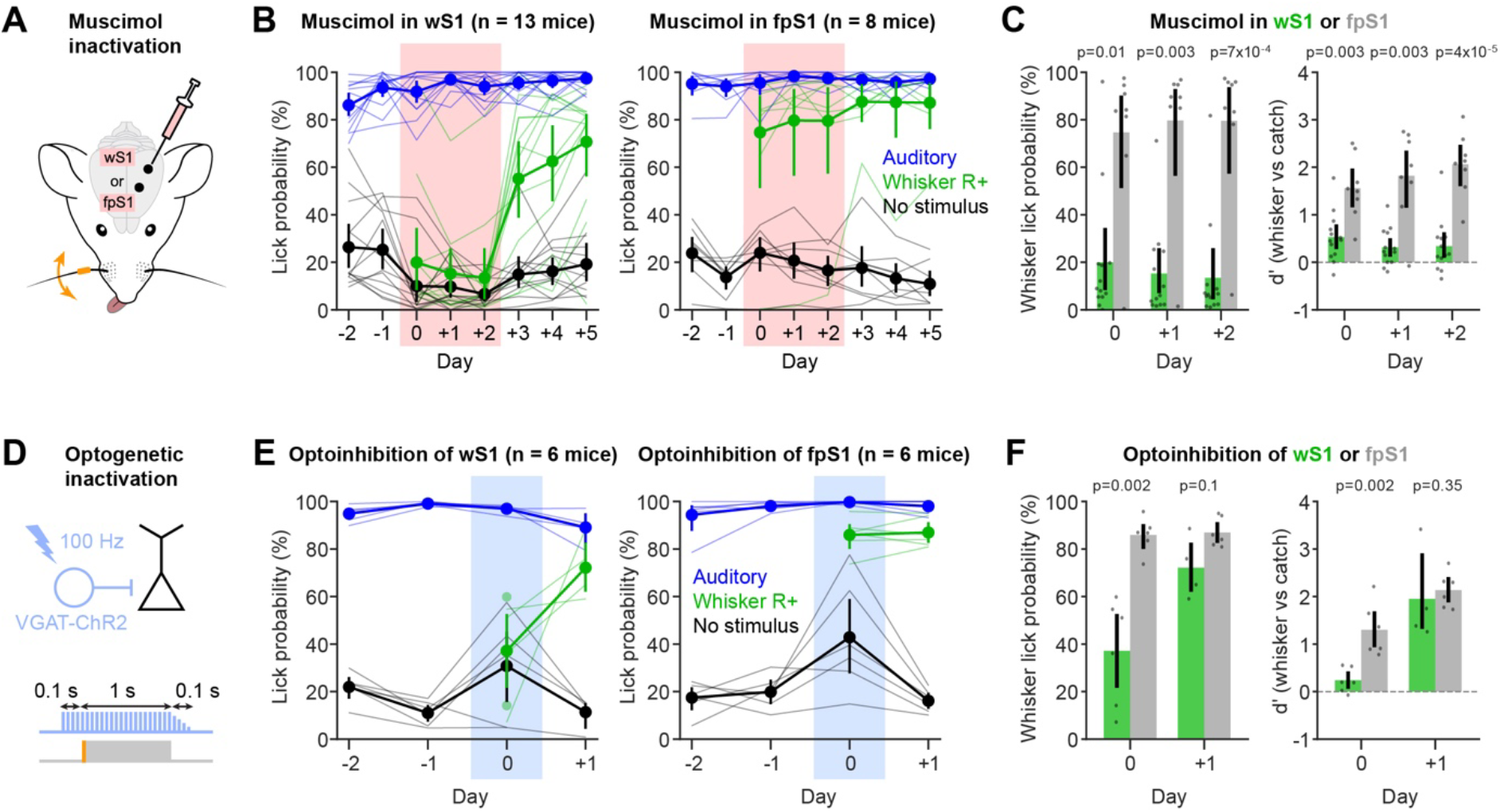
Barrel cortex activity is required for whisker-guided licking during learning. (**A**) Muscimol was injected in the barrel cortex (wS1) or, as a control, in the forepaw primary somatosensory cortex (fpS1) before each session on Days 0, +1 and +2. (**B**) Lick probability on auditory (blue), whisker (green) and catch trials (black) across days, for mice with muscimol inactivation of wS1 (left; n = 13 R+ mice) and fpS1 (right; n = 8 R+ mice). Muscimol days are shaded in red, and recovery days are Days +3 to +5. Thin lines, mice; points, mean ± 95% CI. (**C**) Whisker lick probability (left) and d′ (whisker vs catch trials; right) on the muscimol days, for inactivation of wS1 (green) and fpS1 (grey). Bars, mean ± 95% CI; dots, mice; p-values, Mann-Whitney U test, wS1 vs fpS1, on each day (n = 13 and 8 mice). (**D**) Optogenetic inactivation in VGAT-ChR2 mice. Blue light (100 Hz) was delivered on every trial of Day 0, from 0.1 s before stimulus onset to the end of the 1-s response window, followed by a 0.1 s ramp-down. (**E**) As in panel B, but for optogenetic inactivation of wS1 (left) or fpS1 (right) on Day 0 (blue shading) (Days −2, −1 and 0: n = 6 R+ wS1 mice; n = 6 R+ fpS1 mice) and without light on Day +1 (Day +1: n = 4 R+ wS1 mice; n = 6 R+ fpS1 mice). (**F**) As in panel C, but for optogenetic inactivation on Day 0 and control Day +1.

To further test these findings with a spatiotemporally more precise approach, optogenetic inactivation was performed on Day 0 in VGAT-ChR2 mice (Zhao et al., 2011), in which blue light illumination of the cortical surface activates ChR2-expressing GABAergic interneurons to locally silence excitatory neurons (Guo et al., 2014) (Figure 2D). Optogenetic inactivation was carried out in all auditory, whisker and catch trials, beginning 0.1 s before stimulus onset (or catch trial onset) and lasting throughout the 1-s reporting period. Licking in auditory and whisker trials was rewarded, but not in catch trials. Consistent with the pharmacological results, optogenetic suppression of wS1 activity on Day 0 decreased the probability of licking in response to whisker stimulation compared to mice in which fpS1 was optogenetically inactivated (Mann-Whitney U test, n = 6 R+ wS1-inactivated mice, n = 6 R+ fpS1-inactivated mice: p = 2x10^-3^ on Day 0 for both lick probability and d′) (Figures 2E and 2F). Auditory performance was preserved during optogenetic inactivation of wS1 and fpS1 (Figure 2–figure supplement 1).

These inactivation experiments suggest a specific contribution of neuronal activity in wS1 to drive a licking response to obtain a water reward. Previous experiments inactivating wS1 during closely-related whisker detection tasks performed in expert mice also found important deficits upon pharmacological and optogenetic inactivation of wS1 (Kwon et al., 2016; Le Merre et al., 2018; Miyashita and Feldman, 2013; Oryshchuk et al., 2024; Sachidhanandam et al., 2013; Takahashi et al., 2020, 2016). To test whether wS1 is also required after learning of the current task, we inactivated wS1 or fpS1 with muscimol in a separate cohort of expert R+ mice. Consistent with previous reports, whisker-evoked licking was strongly reduced on the muscimol days in wS1 mice compared with fpS1 mice (muscimol vs control, n = 12 wS1-inactivation mice, n = 6 fpS1-inactivation mice, Mann-Whitney U test: first muscimol day, p = 9x10^-4^; second muscimol day, p = 4x10^-4^) (Figure 2–figure supplement 2). In our inactivation experiments on learning days (Figure 2), it is therefore possible that latent whisker learning could occur but not be expressed through increased licking motor output. We therefore investigated if mice previously inactivated during learning performed better on the first day without inactivation than naive R+ mice on Day 0, but we did not find a significant difference (Figure 2–figure supplement 3). Further experiments in the future specifically designed to investigate the hypothesis of latent learning would be of interest.

### Learning induces a rapid reorganization of barrel cortex population activity

To examine how barrel cortex representations evolve across learning, we performed longitudinal two-photon calcium imaging of the same layer 2/3 neurons expressing GCaMP6f from Day −2 to Day +2 (R+ mice: 3,210 neurons across 19 mice, 169 ± 49 mean neurons per mouse ± SD; R-mice: 2,846 neurons across 16 mice, 177 ± 48 mean neurons per mouse ± SD) (Figure 3 and Figure 3–figure supplement 1A). At the end of each session, including auditory pre-training days (Days −2 and −1), a block of 50 passive whisker stimulations was delivered after task disengagement (three consecutive auditory misses; see Methods) to map sensory responses independently from reward and motor output. Only the last 40 trials of this block were used in neural analyses to ensure that mice had stabilized in a passive state. A striking divergence in the grand-average response emerged where the whisker-evoked responses progressively increased across days in R+ mice but decreased in R-mice (Figures 3A-C). Comparing response amplitudes between pre-learning (Days −2 and −1) and post-learning (Days +1 and +2) periods indicated a significant enhancement of whisker-evoked responses in R+ mice (n = 19 mice, Wilcoxon signed-rank test, p = 3x10^-3^) (Figure 3D) and a significant suppression of whisker-evoked responses in R-mice (n = 16 mice, Wilcoxon signed-rank test, p = 2x10^-3^) (Figure 3E), revealing a reward-dependent bidirectional reshaping of the C2 whisker representation.

**Figure 3.**
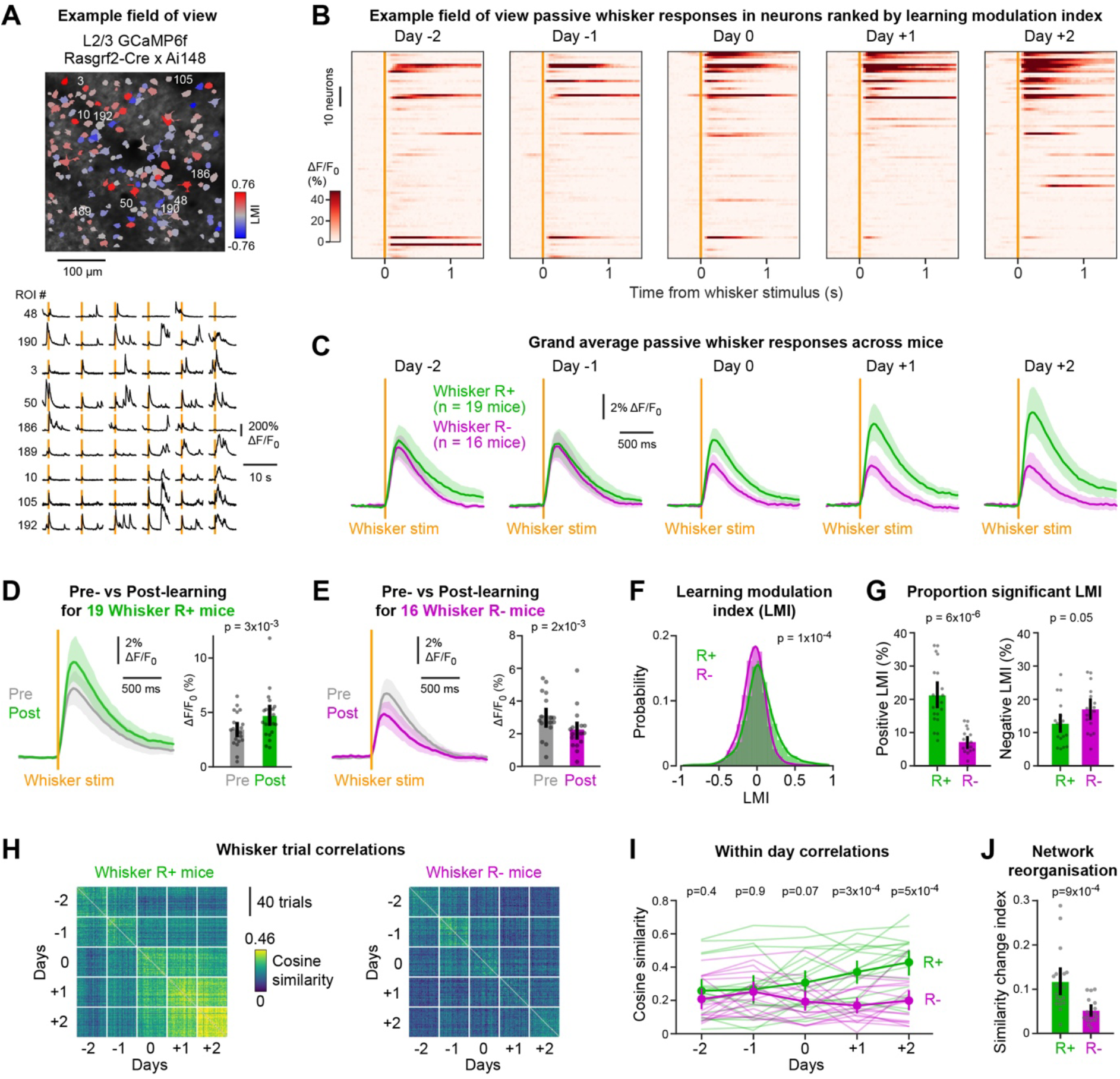
Learning-related reorganization of whisker deflection-evoked barrel cortex population activity. (**A**) Two-photon calcium imaging of barrel cortex layer 2/3 neurons expressing GCaMP6f in Rasgrf2-Cre x Ai148 mice. Top: example field of view of an R+ mouse, with the regions of interest color-coded by the learning modulation index (LMI; red, positive; blue, negative). Bottom: ΔF/F_0_ of example neurons during passive whisker trials. Orange lines, whisker stimulus onset. (**B**) Trial-averaged responses to the passive whisker stimulus on the five imaging days, for the neurons with a significant LMI in the same mouse, ordered by LMI. (**C**) Population responses (ΔF/F_0_, mean over each mouse’s neurons) to the passive whisker stimulus on the five days, for R+ (green; n = 19 mice) and R-mice (magenta; n = 16 mice). Lines, mean ± 95% CI across mice. (**D**) Population whisker response before (Days −2 and −1; Pre, grey) and after learning (Days +1 and +2; Post, green) in R+ mice (n = 19 mice). Left: population responses. Right: response amplitude (mean ΔF/F_0_ over 0-300 ms after stimulus onset). Bars, mean ± 95% CI; dots, mice; p-value, Wilcoxon signed-rank test. (**E**) As in D, for R-mice (n = 16 mice). (**F**) Distribution of the LMI of all imaged neurons in R+ (green; n = 3,210 neurons from 19 mice) and R-mice (magenta; n = 2,846 neurons from 16 mice). The LMI compares each neuron’s responses in the last 40 passive trials of Days −2 and −1 with those of Days +1 and +2 (see Methods). Kolmogorov-Smirnov distance between the two groups, with a p-value from a permutation test where mice were shuffled between groups (10,000 permutations). (**G**) Percentage of neurons with a significantly positive (left) or negative LMI (right) in each mouse, R+ vs R-. Bars, mean ± 95% CI; dots, mice; p-values, Mann-Whitney U test (n = 19 R+, 16 R-mice). (**H**) Cosine similarity between the population responses of all pairs of passive whisker trials (last 40 trials of each day; mean ΔF/F_0_ over 0-300 ms after stimulus onset), averaged across R+ (left; n = 18 mice) and R-mice (right; n = 16 mice). (**I**) Within-day similarity (mean similarity between the different trials of each day) across days. Thin lines, mice; points, mean ± 95% CI; p-value, Mann-Whitney U test, R+ vs R-, on each day (n = 18 R+ and 16 R-mice). (**J**) Network reorganization index: mean of the within-pre-learning (Days −2, −1) and within-post-learning (Days +1, +2) similarity, minus the similarity between pre-and post-learning trials. Bars, mean ± 95% CI; dots, mice; p-value, Mann-Whitney U test (n = 18 R+ and 16 R-mice). In H-J, one R+ mouse with fewer than 40 passive trials on one day was excluded.

A difference in the state of R+ and R-mice could contribute to the apparent changes in whisker-evoked responses during the passive trials. Each behavioral session was terminated when the mouse failed to lick in response to three consecutive auditory trials, which we used as the criterion for task disengagement. The overall motivational state of the R+ and R-mice during the passive whisker stimulation should therefore be similar. Although session duration did not differ comparing R+ and R-mice on learning days, there were differences in the total number of trials and total reward obtained comparing R+ and R-mice (Figure 3–figure supplement 2A–C), which could contribute to differences in the sensory responses. However, for R+ mice, the total reward obtained per session did not change between the pre-and post-learning days (Wilcoxon signed-rank test, p = 0.31), so the enhancement of their responses is not confounded by a change in satiety. Equally, in the passive trials for R+ mice, the whisker angular speed did not change across days (Friedman test, p = 0.80); the tongue rarely protruded in the passive trials; and jaw movements varied across days without following the time course of learning (Figure 3–figure supplement 2D–F). Although we cannot rule out contributions of differing states in R+ and R-mice to the passive whisker response, it seems unlikely to be a major contributor.

We further examined the robustness of our analysis asking if the passive whisker responses were stable or adapted during the 40 trials (Figure 3–figure supplement 3). Population responses were similar in the first and the last 20 trials of the block (R+: p = 0.97, n = 18 mice; R-: p = 0.051, n = 16 mice). Learning-related changes were present in both halves of the block. In R+ mice, responses increased whether the first or the last 20 trials of each day were used (Wilcoxon signed-rank test, first 20 trials, p = 1x10^-3^ and last 20 trials p = 0.016, n = 18 mice). In R-mice, they decreased in the first 20 trials (p = 2x10^-4^, n = 16 mice) and tended to decrease in the last 20 (p = 0.065, n = 16 mice). The changes in whisker sensory-evoked responses comparing across R+ and R-mice were therefore robustly found in the passive trials. However, it is important to note that these learning-related changes were heterogeneous within individual mice at the single-cell level spanning recruitment, enhancement of pre-existing responses, and suppression (Figure 3–figure supplement 1B).

We therefore next identified neurons responsible for this population effect by computing a learning modulation index (LMI) for each cell using ROC analysis between pre-learning (Days −2 and −1) and post-learning days (Days +1 and +2). The LMI distributions differed between R+ and R-mice (two-sample Kolmogorov–Smirnov distance, permutation test, p = 1x10^-4^) (Figure 3F). In every mouse of both groups, the fraction of neurons with a significant LMI exceeded the 5% expected by chance (R+: 19 of 19 mice, Wilcoxon signed-rank test, p = 4x10^-6^; R-: 16 of 16 mice, p = 3x10^-5^) (Figure 3–figure supplement 1C). A significantly larger fraction of neurons acquired a positive LMI (LMI+ neurons) in R+ mice across learning (median 19% vs 6%; n = 19 R+ and 16 R-mice, Mann–Whitney U test, p = 6x10^-6^) (Figure 3G), while the fraction of neurons with a negative LMI (LMI-neurons) tended to be larger in R-mice (median 17% vs 13%; Mann–Whitney U test, p = 0.053) (Figure 3G). As a result, the net fraction of LMI+ minus LMI-neurons was positive in R+ mice (median +8.6%, Wilcoxon signed-rank test against 0, p = 8x10^-3^) and negative in R-mice (median −10.2%, p = 3x10^-3^) (Figure 3–figure supplement 4). Although representational drift in the absence of reward-learning contributes to day-to-day variability of the evoked responses, this drift is symmetrically distributed across LMI+ and LMI-neurons, with no overall change in responsiveness (Figure 3–figure supplement 4).

We next examined how the population representation changed upon learning. We computed trial-by-trial cosine similarity matrices between population activity vectors across all pairs of passive whisker stimulations (Figure 3H). In R+ mice, the within-day similarity increased after learning, while it tended to decrease in R-mice (Mann–Whitney U test R+ vs R-mice; Day 0, p = 0.07; Day +1, p = 3x10^-4^; and Day +2, p = 5x10^-4^; n = 18 R+ and 16 R-mice) (Figure 3I). Trial-to-trial variability (noise power) did not change in either group (Wilcoxon signed-rank test, R+: p = 0.52; R-: p = 0.74). In contrast, response strength (signal power) increased in R+ mice (p = 1x10^-3^) and tended to decrease in R-mice (p = 0.065) (Figure 3–figure supplement 5). Changes in similarity therefore reflect the reliability of whisker responses relative to constant trial-to-trial variability. Across days, R-mice showed a uniform pattern with low similarity, whereas for R+ mice, the average cosine similarity matrix increased with learning. We quantified a network reorganization index, computed as the difference between within-period and between-period cosine similarities, which was significantly larger in R+ than R-mice (Mann–Whitney U test, p = 9x10^-4^) (Figure 3J). Together, these results show that reward-driven learning bidirectionally reshapes the whisker representation of barrel cortex within the first days of whisker-learning.

### Learning-related changes in wS2-and wM1-projecting neurons

Barrel cortex sends whisker information to downstream areas through distinct output pathways. In expert mice performing closely-related whisker detection tasks, task-relevant signals are preferentially routed through the projection to wS2 rather than to wM1 (Vavladeli et al., 2020; Yamashita and Petersen, 2016). To test whether learning-related plasticity differed between these pathways in the current learning paradigm, we retrogradely labelled wS1 neurons projecting to wS2 (wS2p) or to wM1 (wM1p) in a subset of the imaged mice (Figure 4A).

**Figure 4.**
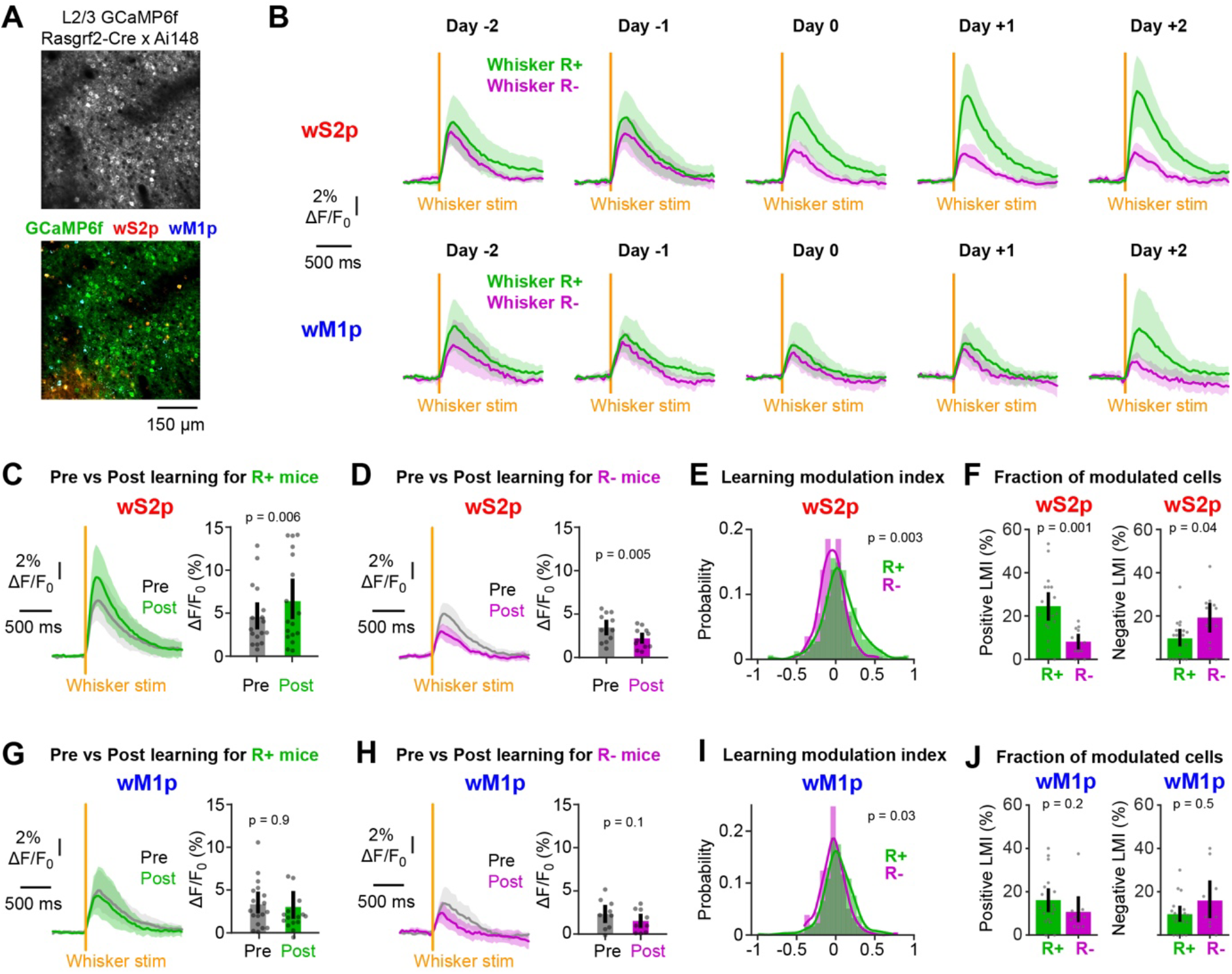
Learning-related change in whisker sensory responses of neurons in wS1 projecting to wS2, but not neurons in wS1 projecting to wM1. (**A**) Top: example field of view of GCaMP6f-expressing neurons. Bottom: the same field of view with the retrogradely labelled neurons: GCaMP6f (green), CTB-labelled neurons projecting to the secondary whisker somatosensory cortex (wS2p, red) and CTB-labelled neurons projecting to the primary whisker motor cortex (wM1p, blue). (**B**) Population responses to the passive whisker stimulus on the five days, for wS2p (top) and wM1p neurons (bottom), in R+ (green) and R-mice (magenta). Lines, mean ± 95% CI across mice (wS2p: n = 17 R+ and 11 R-mice; wM1p: n = 17 R+ and 9 R-mice). In all panels, only mice with at least three neurons of a type are included for analysis. Orange lines, whisker stimulus onset. (**C**) wS2p population whisker response before (Days −2 and −1; Pre, grey) and after learning (Days +1 and +2; Post, green) in R+ mice (n = 17 mice). Left: population responses. Right: response amplitude (mean ΔF/F_0_ over 0-300 ms after stimulus onset). Bars, mean ± 95% CI; dots, mice; p-value Wilcoxon signed-rank test. (**D**) As in panel C, but for R-mice (n = 11 mice) (before learning, grey; after learning, magenta). (**E**) Distribution of LMI for wS2p neurons in R+ (green; n = 328 neurons from 17 mice, 19 ± 10 neurons per mouse, mean ± SD) and R-mice (magenta; n = 253 neurons from 11 mice, 23 ± 11 neurons per mouse). The p-value reports Kolmogorov-Smirnov distance with a permutation test where mice were shuffled between groups (10,000 permutations). (**F**) Percentage of wS2p neurons with a significantly positive (left) or negative LMI (right) in each mouse. Bars, mean ± 95% CI; dots, mice; p-values, Mann-Whitney U test (n = 17 R+ and 11 R-mice). (**G**) As in panel C, but for wM1p neurons in R+ mice (n = 17 mice). (**H**) As in panel D, but for wM1p neurons in R-mice (n = 9 mice). (**I**) As in panel E, but for wM1p neurons (R+, n = 304 neurons from 17 mice, 18 ± 11 neurons per mouse; R-, n = 178 neurons from 9 mice, 20 ± 11 neurons per mouse). (**J**) As in panel F, but for wM1p neurons (n = 17 R+ and 9 R-mice).

wS2p neurons showed similar learning-related changes to the whole population. Their whisker-evoked responses increased in R+ mice (n = 17 mice, Wilcoxon signed-rank test, p = 6x10^-3^) and decreased in R-mice (n = 11 mice, p = 5x10^-3^) (Figures 4B-D). Their LMI distributions differed between R+ and R-mice (Kolmogorov–Smirnov distance, permutation test, p = 3x10^-3^) (Figure 4E). R+ mice had a larger fraction of LMI+ wS2p neurons (Mann-Whitney U test, p = 1x10^-3^) and R-mice a larger fraction of LMI-wS2p neurons (p = 0.035) (Figure 4F). The wM1p neurons showed less change in their whisker-evoked responses across learning. Their response amplitude did not change significantly in either group (R+: n = 17 mice, p = 0.85; R-: n = 9 mice, p = 0.13) (Figures 4B, 4G and 4H). The LMI distributions of wM1p neurons differed between reward groups (p = 0.034) (Figure 4I), but the fractions of LMI+ (p = 0.18) and LMI-(p = 0.48) wM1p neurons did not differ significantly comparing R+ and R-mice (Figure 4J). In R-mice, negative LMI values were larger in magnitude in wS2p than in wM1p neurons (Kolmogorov-Smirnov distance, permutation test, p = 2x10^-3^), while in R+ mice, positive LMI values tended to be larger in wS2p neurons (p = 0.056) (Figure 4–figure supplement 1). Learning-related plasticity was therefore expressed most clearly in neurons projecting to wS2.

### Barrel cortex representations progressively realign during learning

To test whether the reorganization of barrel cortex activity carried information at the single trial level, we trained a logistic regression classifier to discriminate pre-learning (Days −2 and −1) from post-learning (Days +1 and +2) population activity patterns (Figure 5A). Cross-validated decoding accuracy was significantly higher in R+ than R-mice (n = 18 R+ and 16 R-mice; Mann–Whitney U test, p = 1x10^-3^) (Figure 5B). High weights correlated with high LMI and removing the most modulated neurons abolished decoding (one regression slope per mouse, Wilcoxon signed-rank test against 0, p = 1x10^-10^, n = 34 mice) (Figure 5–figure supplement 1). Critically, this decoding approach allowed us to ask whether the transition had already begun on Day 0 by testing the unseen passive trials from that day with the fixed weights of the pre-versus post-learning decoder (Figure 5C). In both groups, the passive whisker-evoked responses on Day 0 could be significantly distinguished from pre-learning and from post-learning days (Wilcoxon signed-rank test, p = 2x10^-5^ for R+ Day 0 vs pre-learning, p = 3x10^-4^ for R+ Day 0 vs post-learning, p = 4x10^-4^ for R-Day 0 vs pre-learning, p = 8x10^-4^ for R-Day 0 vs post-learning). Decoding tended to be more accurate at distinguishing Day 0 from pre-learning than from post-learning (Wilcoxon signed-rank test, p = 0.3 for R+, p = 0.02 for R-; Figure 5D). Together, these results establish that reward-driven learning is rapidly accompanied by a new representational state that emerges during the first whisker learning session.

**Figure 5.**
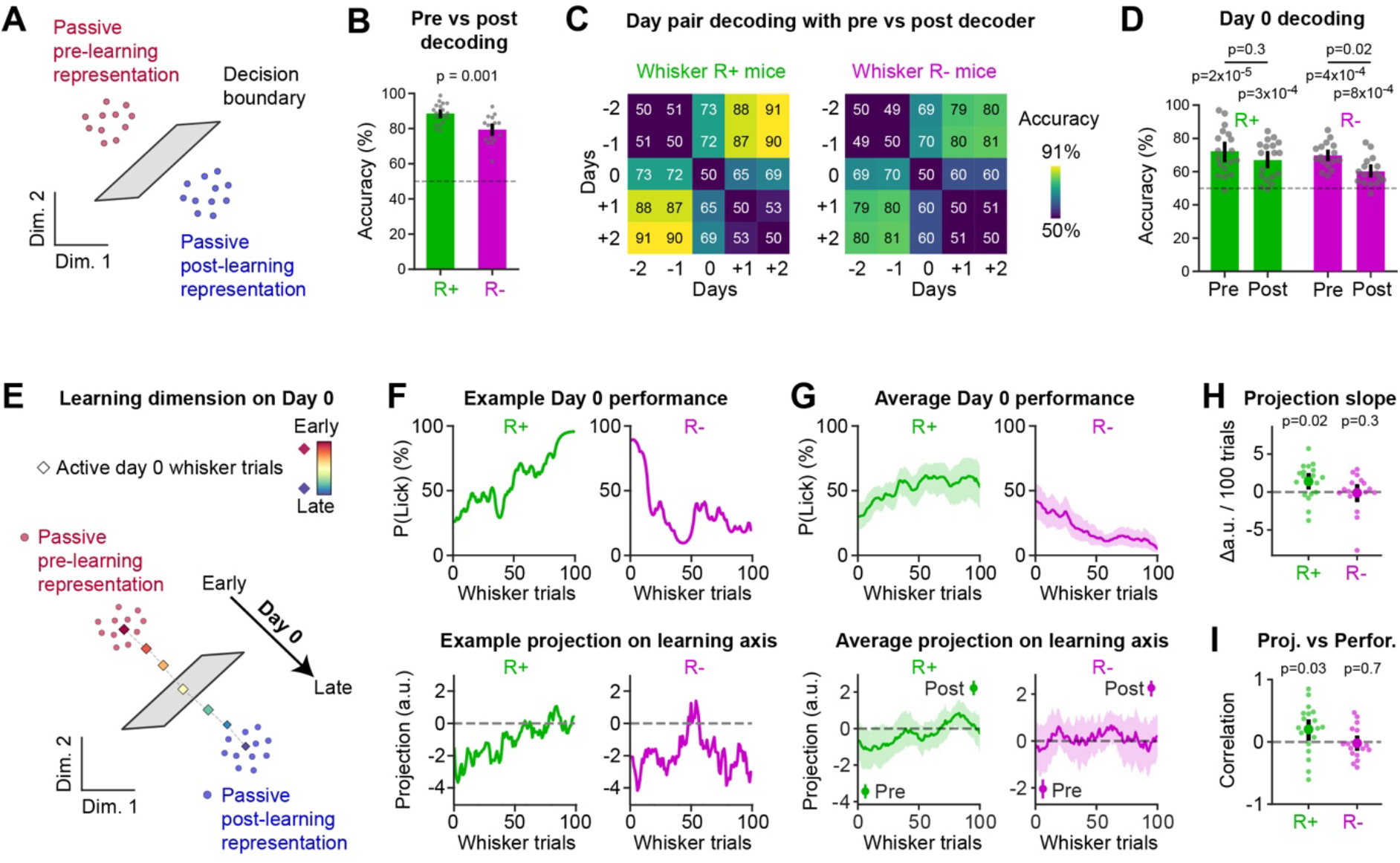
Progressive realignment of barrel cortex representations during learning. (**A**) Schematic of the decoding approach. For each mouse, a logistic regression classifier was trained to discriminate the population responses to passive whisker trials before learning (Days −2 and −1) from those after learning (Days +1 and +2). (**B**) Accuracy of the pre-vs post-learning decoder (10-fold stratified cross-validation), for R+ (n = 18 mice) and R-mice (n = 16 mice). Dashed line, chance level (50%). Bars, mean ± 95% CI; dots, mice; p-value, Mann-Whitney U test. (**C**) Accuracy of the pre-vs post-learning decoder in discriminating each pair of days (held-out trials), averaged across R+ (left) and R-mice (right). (**D**) Decoding accuracy between the pre-learning days and Day 0 (Pre) and between Day 0 and the post-learning days (Post), for R+ and R-mice. Dashed line, chance level. Bars, mean ± 95% CI; dots, mice; lower p-values, Wilcoxon signed-rank tests against chance; upper p-values, Wilcoxon signed-rank tests Pre vs Post within each group (n = 18 R+ and 16 R-mice). (**E**) Schematic of the projection of Day 0 active whisker trials on the learning dimension defined by the decoder. Positive values indicate population activity closer to post-learning responses, negative values activity closer to pre-learning responses; 0 is the decoder boundary. (**F**) Day 0 of an example R+ mouse (left) and R-mouse (right). Top: whisker lick probability. Bottom: projection of the whisker trials on the learning dimension, using activity averaged over 0-180 ms after stimulus onset (before the first lick on nearly all trials), in sliding windows of 10 trials. (**G**) As in panel F, but averaged across mice. Top: lick probability (n = 19 R+ and 16 R-mice). Bottom: projection on the learning dimension (n = 18 R+ and 16 R-mice). Lines, mean ± 95% CI across mice. Pre and Post: projections of the passive pre-and post-learning trials used to train the decoder (mean ± 95% CI across mice). (**H**) Slope of the projection across Day 0 whisker trials. Small dots, mice; large dots, mean ± 95% CI; p-values, one-sided Wilcoxon signed-rank test against 0 (n = 18 R+ and 16 R-mice). (**I**) Pearson correlation between the projection and the lick probability across Day 0 whisker trials. Small dots, mice; large dots, mean ± 95% CI; p-values, one-sided Wilcoxon signed-rank test against 0 (n = 18 R+ and 16 R-mice). One R+ mouse with fewer than 40 passive trials on one day was excluded (see Methods).

Having established that population representations are already partially shifted after Day 0 learning, we next asked whether this transition could be tracked online trial-by-trial during active behavior. To do so, we projected single-trial whisker-evoked population activity from Day 0 onto the decoding axis of the pre-versus post-learning decoder (trained exclusively on the passive whisker trials excluding Day 0) yielding a continuous readout of the shift from naïve to expert representational states (Figure 5E). Day 0 neuronal activity was averaged over 0-180 ms after stimulus onset (i.e. before the first lick on nearly all trials) to minimize potential contributions of lick-related activity to the projection. R+ mice exhibited a progressive increase in both lick probability and projection on this learning dimension across the active Day 0 trials, whereas R-mice showed decreasing lick probability and a flat projection trajectory (Figures 5F and 5G). The linear slope of the projection on the learning dimension over Day 0 trials was significantly positive in R+ mice (one-sided Wilcoxon one-sample test against zero, p = 0.02, n = 18 mice), but not in R-mice (Wilcoxon one-sample test, p = 0.32, n = 16 mice) (Figure 5H). Furthermore, there was significant correlation between the trajectory on the learning dimension and the behavioral learning curve in R+ mice (Wilcoxon one-sample test, p = 0.03 for hit rate and p = 5x10^-3^ for d’), but not in R-mice (Wilcoxon one-sample test, p = 0.7 for hit rate and p = 0.65 for d’) (Figure 5I and Figure 5–figure supplement 2). These results demonstrate that barrel cortex population activity progressively realigns from naïve to expert-like representations during reward learning, tracking behavioral performance trial-by-trial.

### Neurons potentiated by learning preferentially take part in spontaneous reactivations

Spontaneous cortical reactivations of stimulus-evoked ensemble patterns have been shown to occur during inter-trial periods and have been suggested to relate to changes in sensory representations (Miller et al., 2014; Nguyen et al., 2024; Sugden et al., 2020). We therefore examined spontaneous reactivations of the whisker-evoked activity pattern, during the correct-rejection catch trials of the behavioral session where no stimulus is presented (Figures 6A and 6B). Events were detected as peaks of the correlation between population activity and the whisker response template of the same day (see Methods). Comparing the reactivation rates of R+ and R-mice, we found that reward was associated with significantly higher reactivation rates from Day 0 onwards (n = 19 R+ vs 16 R-mice; Mann–Whitney U test, p = 0.010, Day 0; p = 0.035, Day +1; p = 4x10^-3^, Day +2; p ≥ 0.74 before learning) (Figure 6C).

**Figure 6.**
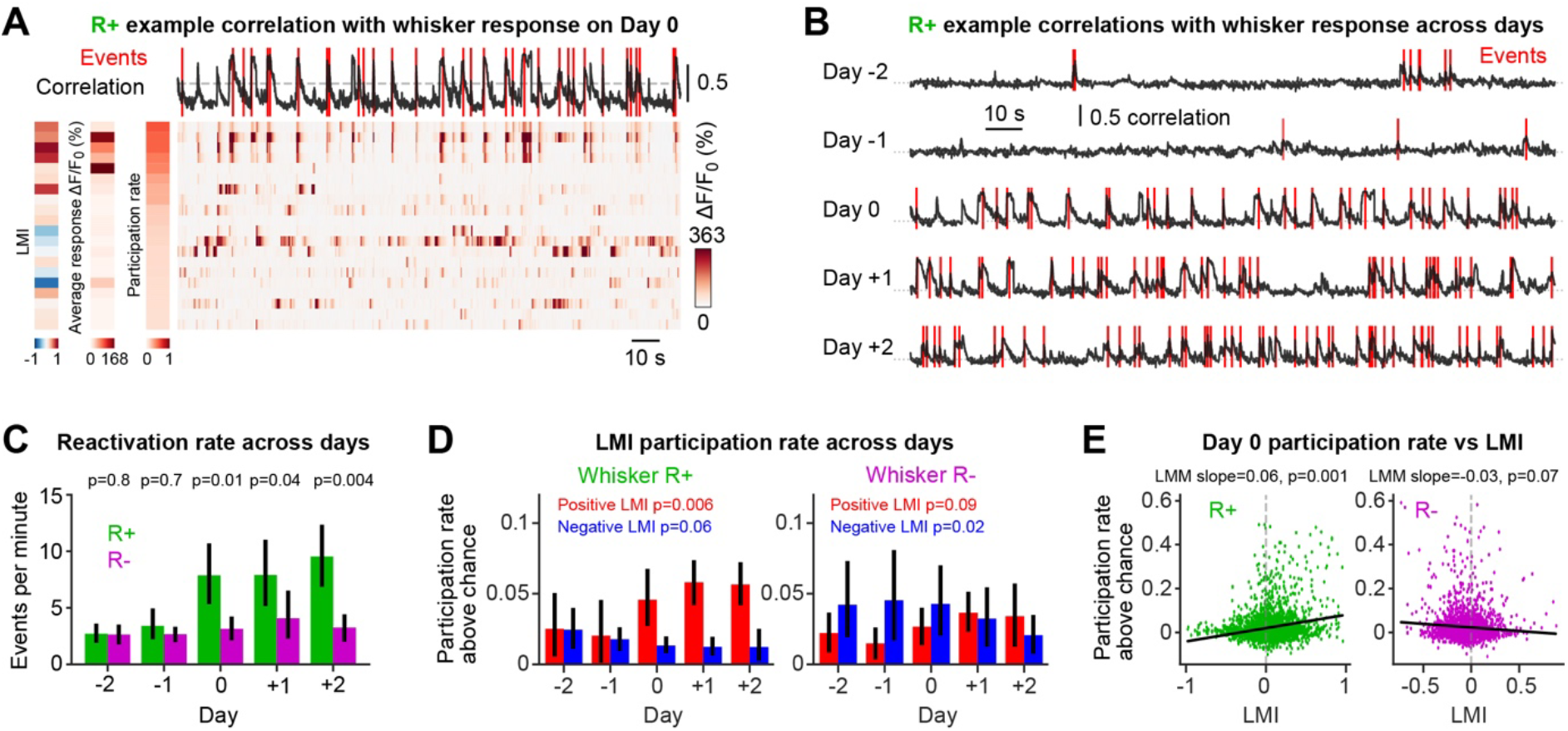
Neurons potentiated across learning are preferentially engaged in spontaneous reactivations. (**A**) Example R+ mouse on Day 0. Right: ΔF/F_0_ of the 20 neurons with the highest participation rate during 180 s of concatenated correct-rejection catch trials. Top: Pearson correlation between the population activity and the whisker response template of the day. Dashed line, detection threshold (99^th^ percentile of circular shift surrogates on the pre-learning days); red lines, detected reactivation events. Left strips, for each neuron: LMI, whisker response template (mean response to the passive whisker stimulus), and participation rate. (**B**) Correlation with the whisker response template on the five days, for the same mouse. Red lines, detected events; dotted lines, zero correlation. (**C**) Reactivation rate (events per minute of correct-rejection trials) across days, in R+ (green; n = 19 mice) and R-mice (magenta; n = 16 mice). Bars, mean ± 95% CI; p-values, Mann-Whitney U test, R+ vs R-, on each day. (**D**) Participation above chance across days of LMI+ (red) and LMI-neurons (blue), averaged per mouse. Bars, mean ± 95% CI across mice; p-values, Wilcoxon signed-rank test against 0 for the slope of the regression of the above chance participation rate against days fitted for each mouse. In panels D and E, only mice with at least three reactivation events on Day 0 were included for analysis (18 R+ mice for LMI+ and LMI-; 13 R-mice for LMI+; and 14 R-mice for LMI-neurons). (**E**) Day 0 participation above chance as a function of LMI, for R+ mice (left; n = 3,104 neurons from 18 mice) and R-mice (right; n = 2,501 neurons from 14 mice). Each dot is one neuron. Participation was measured in held-out neurons: events were detected with a random half of each mouse’s neurons, and participation was measured in the other half (10 random splits). A neuron participated in an event if its ΔF/F_0_ around the event reached 2.5 times its noise SD. Participation above chance is the participation rate minus the rate expected if events occurred at random times. Line, fixed effect of a linear mixed-effects model with a random intercept and a random slope per mouse; slope and p-value (Wald test) above each plot.

Correct-rejection trials contain no licks before trial onset and no licks in the 1-s response window, but mice could lick later in the trial, so we next investigated if licking contributed to event detection. Reactivation rates were nearly constant across the trial, and only slightly higher after the response window, when licks were possible, than in the lick-free part of the trial (R+ mice: median ratio 1.09, Wilcoxon signed-rank test, p = 0.04; R-mice: p = 0.9) (Figure 6–figure supplement 1A). Moreover, when events were detected using only the lick-free part of each trial (−1 to +1 s), with detection thresholds recomputed from this lick-free activity, reactivation rates on Day 0 remained higher in R+ than in R-mice (Mann-Whitney U test, p = 0.029) (Figure 6–figure supplement 3A). We also checked that reactivations did not reflect global coactivation but were specific to the whisker response by detecting events with templates in which neuron identities were shuffled. Shuffled templates detected significantly fewer events than the real template, with the same activity and detection threshold, in both groups and on every day (Wilcoxon signed-rank test of the per-mouse difference, p ≤ 5x10^-3^) (Figure 6–figure supplement 1B). The increased reactivation rate in R+ mice compared to R-mice therefore seems to be robust across various analyses.

Given the differences between reward groups, we wondered if the outcome of the preceding trial influenced reactivation rates. However, we did not find that Day 0 reactivation rates depended on the outcome of the previous trial (rewarded or unrewarded; R+: p = 0.62; R-: p = 0.28) or on the outcome of the last whisker trial (hit or miss; R+: p = 0.47; R-: p = 0.064). Equally, we found that, R+ mice still had higher reactivation rates than R-mice after unrewarded trials (Mann-Whitney U test, p = 9x10^-3^) (Figure 6–figure supplement 1C). In future experiments and analyses, it will therefore be important to specifically focus on which aspects of reward learning are important for driving the increased reactivation rate in R+ mice.

We next computed how much each neuron took part in these reactivations. To avoid measuring a neuron’s participation in events it helped to detect, events were detected with a random half of the neurons and participation was measured in the other half (see Methods). Because frequently active neurons cross the participation threshold more often by chance, we analyzed participation above the level expected from each neuron’s activity on the same day. In R+ mice, the participation of significantly LMI+ neurons increased across days, starting on Day 0 (per-mouse slope across days, Wilcoxon signed-rank test, p = 6x10^-3^, n = 18 mice), whereas that of LMI-neurons did not increase (p = 0.06) (Figure 6D). In R-mice, the participation of LMI-neurons decreased across days (p = 0.025, n = 14 mice), whereas that of LMI+ neurons did not change significantly (p = 0.09, n = 13 mice). Participation thus followed the dominant direction of plasticity in each group. Consistently, Day 0 participation increased with LMI in R+ mice (linear mixed-effects model with mouse as random intercept and slope, p = 1x10^-3^, n = 18 mice) and tended to decrease with LMI in R-mice (p = 0.07, n = 14 mice) (Figure 6E). These results for R+ mice held as well with raw participation rates without the chance correction (Figure 6–figure supplement 2), at other detection and participation thresholds, and restricted to the lick-free part of the trial (Figure 6–figure supplement 3). Together, these results indicate that neurons whose sensory responses were strengthened by reward-driven learning were preferentially engaged in spontaneous reactivation events.

## Discussion

Here, we developed a task in which mice formed a whisker-to-lick sensorimotor association within a single behavioral session (Figure 1). Pharmacological and optogenetic inactivation suggested an important role for neuronal activity in wS1 (Figure 2). Neuronal activity in wS1 exhibited a reorganization of whisker sensory representations across reward learning (Figure 3), with neurons projecting to wS2, but not neurons projecting to wM1, exhibiting significant learning-related changes (Figure 4). The changes in whisker representation in wS1 began on the first day of whisker learning, and already during this first single session representational change correlated online with task learning (Figure 5). Finally, we found that neurons whose stimulus responsiveness was strengthened by learning were preferentially engaged in spontaneous reactivation events (Figure 6). Future experiments will be needed to test causal mechanisms.

### Rapid whisker-dependent reward-based goal-directed sensorimotor learning

We aimed to design a task to study rapid whisker-dependent reward-based goal-directed sensorimotor learning (Figure 1). Mice learned through trial-and-error that the whisker stimulus was predictive of reward availability upon licking for R+ mice, but not for R-mice. In our task the mice were pretrained to lick in response to an auditory stimulus, so the overall knowledge that a brief sensory stimulus should drive a licking response for reward had already been learned by mice before the whisker stimulus was introduced in the active behavioral sessions on Day 0. Our goal was to specifically isolate what is learned by the mice to the precise association of a millisecond timescale impulse to a single whisker stimulus predicting reward availability (R+ mice) or not (R-mice) upon execution of a well-known action (licking). Even in the first whisker trials, both R+ and R-mice licked above the false alarm rate, suggesting a generalization of the stimulus-action coupling rule learned for auditory stimuli and generalized to the whisker stimulus. Subsequently, as mice gathered evidence across trials, licking probability in response to the whisker stimulation increased for R+ mice and decreased for R-mice. Learning therefore might be equally important in both cohorts of mice. R+ mice learn to increase licking probability for whisker stimulation, whereas R-mice learn to withhold licking in response to whisker stimuli. At the population level the performance of R+ and R-mice diverged after 22 whisker trials over 14 minutes, suggesting that reward evidence is accumulated across trials.

It is also interesting to note that on the whisker learning day, the whisker stimulus is not entirely a novel stimulus for the mice. They have received the same whisker stimulus at the end of each of the previous auditory training days in an entirely passive context. So, in part, the learning on Day 0 might also involve unlearning of a whisker stimulus as an uninteresting sensory input delivered in a sated state and converting it into an important signal predictive of reward.

### Causal roles of neuronal activity in wS1

In order to examine potential causal roles for neuronal activity in wS1, we carried out inactivation experiments (Figure 2). We found that R+ mice failed to increase licking probability in response to whisker stimulation when wS1 was inactivated either pharmacologically or optogenetically. The inactivation experiments exhibited spatial specificity since inactivation of fpS1 did not impair learning and were also modality-specific since the auditory performance remained high during the inactivation. We did not find evidence for latent learning during the inactivation experiments, but carefully designed experiments must specifically address this point in the future. Given that mice without wS1 can learn whisker detection tasks in other experimental paradigms (Hong et al., 2018; Hutson and Masterton, 1986), there must be other brain regions that can contribute to learning.

Although these inactivation experiments reveal the necessity of neuronal activity in wS1 to increase licking probability in response to the whisker stimulus in our task, they do not necessarily suggest that wS1 is a locus of learning or synaptic plasticity. Indeed, upon inactivation of wS1, expert mice also decreased their whisker performance, suggesting that wS1 is necessary for task execution. Future experiments, therefore, need to address whether learning-related changes in wS1 are essential for task learning. One important avenue of future research could be to test if learning occurs in mice in which synaptic plasticity in wS1 has been blocked, for example by inhibiting calcium/calmodulin-dependent protein kinase II, which is often required for the induction of long-term potentiation (Hardingham et al., 2003; Majumder et al., 2026).

### Representational change of wS1 neuronal activity across days

By longitudinal imaging of GCaMP6f expressed in layer 2/3 neurons of wS1 during passive whisker stimuli delivered after mice had stopped licking in response to auditory stimuli, we found that neurons on average increased their response to whisker stimulation in R+ mice whereas the whisker response decreased in R-mice (Figure 3). Differentiating between two largely non-overlapping classes of projection neurons, we found that neurons located in wS1 projecting to wS2 exhibited overall increases in the whisker-evoked responses in R+ mice and decreases in R-mice, whereas wS1 neurons projecting to wM1 did not show prominent changes in whisker-evoked responses in either R+ or R-mice (Figure 4). These projection-specific changes are consistent with the preferential routing of task-relevant information through the wS1-to-wS2 pathway reported previously in expert mice performing closely-related whisker detection tasks (Kwon et al., 2016; Vavladeli et al., 2020; Yamashita and Petersen, 2016).

A potential confound in comparing the passive whisker-evoked responses in the R+ mice and R-mice is that they may not be in the same state. The passive whisker stimuli were delivered after mice stopped licking in 3 consecutive auditory trials, which indicated disengagement in the task, presumably due to satiety. Comparing across R+ and R-mice, we found that session duration did not differ, but there were differences in the total number of trials and total reward obtained. Within R+ mice, we found little change in these task measures across learning, and in addition we found no differences in whisker, tongue or jaw movements across learning. Therefore, although we cannot rule out contributions of different states to changes in neuronal representations, it seems unlikely to be a major factor. Further experiments attempting to control for various differences in behavioral states will be necessary to address this question more directly.

It is also important to note that many neurons in each mouse increased in responsiveness and many neurons decreased in responsiveness, and thus within individual mice there is a large spread in the learning modulation index. Indeed, changes in responsiveness in the absence of reward-based learning, often termed representational drift, appears prominent in various cortical regions across sensory modalities, including barrel cortex (Ahmed et al., 2024; Bauer et al., 2024; Marks and Goard, 2021; Nguyen et al., 2024; Wang et al., 2022). Here, we also noted changes in whisker-evoked responses across the baseline days, which presumably contribute to our measure of learning-associated representational change. However, the baseline representational drift was equally distributed across increasing versus decreasing responses to whisker stimulation, whereas our learning modulation index changed differently in R+ and R-mice showing increases and decreases, respectively, in overall responsiveness. At the level of any given individual neuron, therefore, we cannot state that the changes were learning-related and not due to representational drift, but at the population level we can distinguish these through the asymmetrical change according to reward group.

### Representational change of wS1 neuronal activity within a single-session

By designing a task that could be learned in a single session, we could track the representational transition as it unfolded trial-by-trial in the first whisker learning session (Figure 5). We defined a learning dimension from the ability of a population decoder to distinguish the representational change in passive whisker responses comparing pre-learning days (Days −1 and −2) with post-learning days (Days +1 and +2). Projecting neuronal activity trial-by-trial on the first day of whisker learning, as the mice learned the reward contingency, revealed that the representation of the whisker stimulus gradually changed from being more aligned to pre-learning day representation to being more aligned with the post-learning representation in a manner correlated with behavioral performance for R+ mice. Importantly the decoder was constructed without including any data from the first day of whisker learning. This goes beyond prior work correlating neuronal selectivity with performance across days (Banerjee et al., 2020; Chéreau et al., 2020; Goltstein et al., 2013; Henschke et al., 2020; Kato et al., 2015; Makino and Komiyama, 2015; Peron et al., 2015b; Poort et al., 2015) and shows that representational changes are concurrent with behavioral acquisition, rather than a delayed consequence. In contrast, a recent study in the auditory cortex also found that sensory cortex contributed to rapid acquisition of task contingencies within tens of trials, but rapid learning in that study did not enhance stimulus representations, but rather drove late-in-trial reward-prediction signals (Drieu et al., 2025). In the future, it will be interesting to examine if differences in task design or divergent sensorimotor processing across different sensory cortices might account for the differences in reward-related neuronal plasticity of somatosensory versus auditory cortices.

### Spontaneous reactivations hof whisker deflection-evoked neuronal activity

Investigating periods without sensory stimuli or licking on whisker learning days, we found a high rate of spontaneous activity events resembling the passive whisker stimulus-evoked representation in R+ mice, but not R-mice (Figure 6). The analysis was robust to multiple variations, including taking account of spontaneous firing rates, differing thresholds for event identification and neuronal participation. Most importantly, the neurons participating in spontaneous reactivations were identified in held out analyses, in which only half of the neurons were used to identify events, and the other half of the neurons were used to characterize the properties of the co-active ensemble. Altogether, these analyses indicated robust reactivation of whisker-responsive neurons during spontaneous intertrial periods in R+ mice, and, interestingly, neurons gaining responsiveness across reward learning (LMI+ neurons) preferentially took part in reactivations in R+ mice.

In previous work studying the visual association cortex of mice learning a visual drifting-grating orientation discrimination task, reactivations of important visual sensory cues also occurred prominently in rest periods after learning sessions (Sugden et al., 2020). Furthermore, neuronal ensemble reactivations have been suggested to predict the direction of change of future visual representations even during passive stimulus viewing, suggesting that they could act as an upstream selection process rather than a faithful replay of past activity (Miller et al., 2014; Nguyen et al., 2024). Our analyses do not distinguish whether the spontaneous reactivation events are a result or a cause of learning. Indeed, the two concepts might be intertwined in a positive feedback loop. If we imagine a form of Hebbian “fire-together-wire-together” plasticity (Hebb, 1949), then one might expect a strengthening of synaptic connectivity between relevant co-active excitatory neurons across learning in R+ mice. In turn, the strengthened synaptic connections might help make these neurons spontaneously co-active through reciprocal excitation, and that very same co-activity might help further reinforce their synaptic connectivity strength through further Hebbian learning, in a repeating self-reinforcing cycle. In future experiments, it will be of great interest to specifically perturb spontaneous reactivations to examine whether they play causal roles in learning in our task.

In summary, here, we found that reward-based, goal-directed sensorimotor learning was correlated with changed neuronal representations of a relevant sensory stimulus in sensory cortex through: i) enhanced whisker responses in R+ versus R-mice; ii) differing contributions according to the neuronal projection type; and iii) increased spontaneous reactivation rates of ensembles representing the rewarded sensory stimulus. Future experiments must investigate potential causal roles for our current correlational observations.

## Methods

### Animals

All procedures were approved by the Swiss Federal Veterinary Office (License numbers VD-1628 and VD-3753) and were conducted in accordance with the Swiss guidelines for the use of research animals. We used both male and female adult C57BL/6 mice of at least 5 weeks at the time of surgery. For calcium imaging experiments we used Rasgrf2-dCre mice [B6;129S-Rasgrf2 <tm1(cre/folA)Hze>/J, JAX: 022864] (Harris et al., 2014) crossed with Ai148 mice [B6;Ai148(TIT2L-GC6f-ICL-tTA2)-D, JAX: 030328] (Daigle et al., 2018), which express the green fluorescent calcium indicator GCaMP6f in layer 2/3 neurons. For optogenetic stimulation of inhibitory neurons, we used VGAT-ChR2 mice [B6.Cg-Tg(Slc32a1-COP4*H134R/EYFP)8Gfng/J, JAX: 014548] (Zhao et al., 2011). For pharmacological inactivation experiments we used wild-type mice.

The imaging dataset comprised 35 mice (19 R+ and 16 R-). The inactivation experiments used three cohorts: muscimol inactivation during learning (13 wS1 and 8 fpS1 mice), optogenetic inactivation during learning (6 wS1 and 6 fpS1 mice) and muscimol inactivation in expert mice (12 wS1 and 6 fpS1 mice).

### Head plate surgery and intrinsic optical imaging

Mice were first implanted with a metal head-post under anesthesia (1.5-2.5 % isoflurane). Before the start of surgery, carprofen was injected (7.5 mg/kg at 1.5 mg/ml, subcutaneous) for general analgesia, and a mix of lidocaine and bupivacaine below the scalp as local analgesic. An ocular ointment (Vita-Pos, Pharma Medica AG, Switzerland) applied over the eyes prevented drying during the surgery. A solution of povidone-iodine (Betadine, Mundipharma Medical Company, Bermuda) was applied for skin disinfection before incision. We monitored body temperature throughout the surgery and it was maintained at 37°C with a heating pad (ThermoStar, Intellibio, France). After excising the scalp with surgical scissors, the exposed skull was cleaned with cotton buds and a scalpel blade to remove the periosteal tissue. After further disinfection with Betadine, the skull was dried with cotton buds and made optically clear using a thin layer of super glue applied over the dorsal part of the skull (Loctite super glue 401, Henkel, Germany). We then glued a custom-made head fixation plate above the right hemisphere and further secured it with self-curing denture acrylic (Paladur, Kulzer, Germany or Ortho-Jet, Lang, USA). Finally, a second thin layer of the glue was applied homogeneously on the left hemisphere to ensure a smooth and equally transparent field of view. This transparent skull preparation was used to perform intrinsic optical signal (IOS) imaging. Mice were returned to their home cages and ibuprofen (Algifor Dolo Junior, Verfora SA, Switzerland) was added to the drinking water for three days after surgery.

Following head plate implantation, IOS imaging was performed on the left hemisphere in order to localize the right C2 whisker representation in wS1 and wS2, as well as the location of fpS1. Animals were lightly anesthetized (isoflurane at 1%) and head-fixed under a CMOS camera coupled to a stereomicroscope (Leica MZ9.5). We acquired a reference image of the surface vasculature under green light (525 nm, Thorlabs LED). Subsequently, under red light (630 nm, Thorlabs LED), the right C2 whisker or the right paw was stimulated by a capillary glass glued to a piezoelectric actuator (PICMA, PI Ceramic). Each trial consisted of 4 s of rectangular deflections at a 10 Hz frequency. We interleaved stimulation with no-stimulation trials and the average activation map was computed using a custom MATLAB-based software. The resulting map was then overlaid on the anatomical image to locate the region of interest relative to surrounding blood vessels.

### Cranial window surgery for two-photon imaging and retrograde labelling

For mice used for two-photon calcium imaging, a circular craniotomy with a 4 mm diameter was performed over the C2 column of wS1 identified with IOS. After the bone had been removed, the surface of the cortex was rinsed with Ringer solution and the dura was removed using a 27G sharp needle bent as a hook. To retrogradely label wS1 neurons projecting to wS2 and wM1, cholera toxin subunit-B (CTB) conjugated with two different Alexa-Fluor dyes (Molecular Probes, Invitrogen; 100 nL, 0.5%, wt/vol) were injected in these areas. A thin glass pipette (PCR Micropipettes 1-10 µL, Drummond Scientific Company, USA) was first pulled and then the tip was broken using a tissue to obtain a 21-27 µm inner tip diameter. The pipette was filled with mineral oil and then tip-filled with the CTB solution. Two injections in the cortex at depths of 300 µm and 500 µm below the pia were performed of 50 nL at each depth for wS2 and 100 nL for wM1 (at stereotaxic coordinates 1 mm anterior and 1 mm lateral from bregma). Injections were performed using a single-axis oil hydraulic micromanipulator (Narishige, Japan) at 20 nL per minute with 5 min of waiting time after injection. A 4 mm glass window was then fixed over wS1 using super glue and self-curing denture acrylic.

### Behavioral task and training

We trained head-fixed, water-restricted mice to perform a sensory detection task. During the behavioral experiments, all right whiskers were trimmed except for the C2 whisker, and the mice were water restricted to 0.9-1.2 mL of water/day (depending on initial body weight). Mice were trained daily with one session per day; their weight and general health status were carefully monitored using score sheets. Mice were habituated to be head-fixed to a metal post with their head held at a 30-degree angle. During the first session, mice were also habituated to licking from a waterspout positioned on their right side which delivered 5 µL of water in response to some licks. Next, we trained them in an auditory detection task for 5-7 days during which licking in a 1 sec response window after the auditory cue (10 ms, 10 kHz tone of 74 dB embedded in the continuous background white noise of 80 dB) was rewarded. Inter-trial interval was uniformly sampled between 6 s and 10 s including a final 2 s to 5 s no-lick window during which mice were required not to lick for the trial to be initiated. When two consecutive days of stable expert performance (hit rate ≥ 80% and false alarm rate ≤ 30%) in the auditory detection task was reached (Days −2 and −1), whisker detection training was introduced for the next three days (Days 0, +1 and +2). Whisker deflection was achieved with a 30 mT magnetic pulse driven by a 3 ms cosine impulse sent to an electromagnetic coil positioned under the mouse and acting on a small iron particle placed on the right C2 whisker. Trials in auditory and whisker training days were presented pseudo-randomly in blocks of 20 trials. Blocks consisted of 10 auditory and 10 catch (no stimulus) trials on auditory training days and 7 whisker, 3 auditory and 10 catch on whisker training days. If mice licked the reward spout within the reward time window of 1 s from stimulus onset, it was considered a hit trial, and they obtained a 5 µL drop of water. If they did not lick within the reward window, it was considered a miss trial, and no reward was delivered. Each mouse was randomly assigned to the rewarded (R+) or non-rewarded (R-) group at the start of the training protocol. While auditory trials were rewarded for both groups throughout training, only R+ mice obtained a water reward in whisker trials, whereas R-mice did not obtain a water reward if they licked in response to whisker stimulation. Catch trials were never rewarded. Task trials were analyzed up to task disengagement, defined for each session as three consecutive auditory misses.

In every two-photon imaging session (Days −2 to +2), we additionally presented 50 passive whisker stimuli at the end of the session, after the mouse had disengaged from the task (three consecutive missed auditory trials). The lick spout was kept in place. Passive stimulation continued until 50 whisker stimuli without a lick in the response window had been delivered, and the last 40 trials were analyzed. Analyses of passive responses used the last 40 passive trials of each day, so that mice could settle into the passive state. In two sessions, fewer than 50 passive stimuli were erroneously delivered (42 and 33).

In a subset of R+ mice, we verified that licking depended on the whisker stimulus itself rather than a potential artefactual noise linked to the delivery of the whisker stimulus: one day after whisker training, a particle test session was performed where the iron particle was initially placed on the whisker for a first block, then removed from the whisker during a block of the session and then put back (particle test; Figure 1–figure supplement 1C).

### Optogenetic inactivation

Optogenetic inactivation of cortical excitatory neurons was achieved indirectly by activating ChR2-expressing GABAergic interneurons in VGAT-ChR2 mice (Zhao et al., 2011). Blue light (470 nm; MBL-F473/200 mW, Changchun New Industries Optoelectronics Technology, China) was delivered through a 200 µm, 0.22 NA patch cable (M84L01, Thorlabs, USA) fitted with a ceramic ferrule cannula (CFMXA05, Thorlabs, USA). The target area, wS1 or fpS1, was identified by IOS imaging and the skull was thinned above the site to facilitate light penetration. The remainder of the exposed skull was covered with Kwik-Cast to prevent off-target illumination of adjacent cortical areas. A blue masking light (M490L4, Thorlabs, USA) was placed in the animal’s visual field throughout inactivation sessions to prevent the light stimulus from serving as a visual cue. Light stimulation consisted of a 100 Hz pulse train with a 50% duty cycle, delivered on all trials (whisker, auditory and catch). Stimulation onset was 100 ms before stimulus presentation and lasted 1 s to cover the full reward window, followed by a 100 ms linear ramp-down to prevent rebound excitation. Light power was measured at 10 mW at the fiber tip. Inactivation was performed on Day 0, during initial whisker learning, and most mice were trained for one additional session without light stimulation on Day +1 (4 of the 6 wS1 mice and all 6 fpS1 mice).

### Pharmacological inactivation

Cortical inactivation was achieved by injection of the GABA-A receptor agonist muscimol (5 mM in Ringer solution; Biotrend, USA). A small craniotomy was drilled above the C2 barrel column of wS1 or above fpS1 identified by IOS imaging. Muscimol was injected through a glass pipette of 21-27 µm in diameter at a rate of 20 nL/min, to a total volume of 400 nL distributed across four depths ranging from 200 to 800 µm below the pia. Injections were performed 45-60 min before the start of the behavioral session. Inactivation was performed during the three days of whisker training (Days 0, +1, +2); mice were then trained for three additional recovery days without injection (Days +3, +4, +5).

To test the role of wS1 after learning, a separate cohort of mice was trained in the whisker task until expert performance. After two baseline sessions at stable expert performance, muscimol was injected in wS1 or fpS1 before the next session, Ringer solution was injected in the same way before the following session, and muscimol was injected again before the session after that (Figure 2–figure supplement 2). Injection volume and depths were as above.

### Two-photon imaging

A custom made two-photon microscope was used to perform calcium imaging experiments described in this paper. The microscope was equipped with a galvo-resonant mirror pair (8 kHz CRS, Cambridge Technology, USA), allowing a frame rate of 30 Hz for a resolution of 512 × 512 pixels. A femtosecond tunable infrared laser (Mai Tai or InSight DeepSee, Spectra Physics–Newport, USA) was fed into the light path at a wavelength of 940 nm to excite the genetically-encoded calcium indicator GCaMP6f. Light emission was detected with a GaAsP photosensor module (H10770PA-40, Hamamatsu, Japan), and signal acquisition was performed with National Instruments hardware (NI PXIe-1073, NI PXIe-6341, National Instruments, USA). The microscope head was movable and controlled in three dimensions by motors (Luigs and Neumann, Germany). A 16x immersion objective (16x Nikon CFI LWD, Japan) was used for all the imaging. The system was operated by the Matlab-based software ScanImage (Pologruto et al., 2003) (Vidrio Technologies, USA). For each mouse, the same field of view was imaged for five consecutive days (Days −2 to +2).

### Movement tracking

Whisker and orofacial movements were filmed with a high-speed camera (CL 600 X 2/M, Optronis, Germany) running at a frame rate of 200 Hz, an exposure time of 4 ms and a resolution of 128x128 pixels in 11 R+ mice and tracked with DeepLabCut (Mathis et al., 2018). The whisker angle was computed from the base and the tip of the C2 whisker, and the whisker angular speed as the absolute value of its time derivative (deg/sec); tongue and jaw speeds were the frame-to-frame displacements of the tongue and the jaw (pixels/sec). For each mouse and day, speeds were averaged over the last 40 passive trials and over the 1 sec after stimulus onset (Figure 3–figure supplement 2D-F).

### Quantification and data analysis

#### Behavioral performance

Lick probability was computed for each session and trial type as the fraction of trials with a lick in the 1-s response window. Sensitivity was quantified as d′ = Z(hit rate) -Z(false-alarm rate), where Z is the inverse of the standard normal cumulative distribution function, with whisker trials as signal and catch trials as noise. For the analyses of blocks of trials (Figure 2–figure supplement 3), trials were grouped into consecutive blocks of 20 trials of all types, and lick probability and d′ were computed within each block in the same way.

To obtain a continuous trial-by-trial estimate of behavioral performance rather than averaging binary hit/miss outcomes across mice, lick probability was estimated trial-by-trial using a Bayesian state-space model implemented with PyMC (Smith et al., 2004). For each session and stimulus type, binary hit/miss outcomes were modelled as Bernoulli observations with a lick probability that evolved as a Gaussian random walk in logit space. The logit-transformed lick probability at each trial t was modelled as logit(p_t_) = x_t_, where x_t_ followed a Gaussian random walk. Posterior inference was performed by MCMC sampling, the posterior mean of p_t_ was taken as the estimated lick probability at each trial, and the 80% credible interval was used to characterize uncertainty.

#### Two-photon calcium imaging preprocessing

To extract time-varying somatic calcium signals, we used the Suite2p toolbox (Pachitariu et al., 2017). For each mouse, the recordings of the five imaging days were concatenated and processed together, and motion correction registered all sessions to a common reference. Each region of interest (ROI) was detected once and its fluorescence extracted from the same pixels on all five days. Neuropil contamination was corrected by subtracting the fluorescent signal from a surrounding ring F_neuropil_(t) from somatic fluorescence: F(t) = F_soma_(t) -0.7 × F_neuropil_(t). The baseline F_0_(t) was computed with a maximin filter: the fluorescence was low-pass filtered at 1 Hz, then a rolling minimum followed by a rolling maximum was applied over a 60-s window, and the result was smoothed with a Gaussian kernel (σ = 5 s). ΔF/F_0_(t) was computed as (F(t) -F_0__F(t)) / F_0__soma(t), where F_0__F(t) and F_0__soma(t) are the baselines of the neuropil-corrected and of the raw somatic fluorescence.

Labelling of the wS2-and wM1-projecting neurons was performed by registering the mean red and far red CTB images with the mean GCaMP6f image with the Python-based SimpleElastix toolbox (https://simpleelastix.github.io/). Every ROI was then classified by visual inspection of its average fluorescence into wS2-projection, wM1-projection or non-labelled with a custom Python GUI.

For analysis, ΔF/F_0_ was aligned to the onset of each trial at the nominal frame rate of 30 Hz, from 2 sec before to 6 sec after onset; the frame nearest to stimulus onset was taken as time 0. Unless stated otherwise, the mean ΔF/F_0_ from 1 s before onset to onset was subtracted from each trial (baseline), and the response of a neuron to a stimulus was its mean baseline-subtracted ΔF/F_0_ from 0 to 300 ms after stimulus onset for passive trials and from 0 to 180 ms for active trials.

#### Learning modulation index

To quantify learning-induced changes in whisker-evoked responses at the single-cell level, a learning modulation index (LMI) was computed for each neuron using receiver operating characteristic (ROC) analysis. For each cell, the response to each of the last 40 passive whisker trials of each day was computed (mean baseline-subtracted ΔF/F_0_ over 0-300 ms after stimulus onset). ROC analysis was then performed by treating pre-learning trials (Days −2 and −1 pooled) as class 0 and post-learning trials (Days +1 and +2 pooled) as class 1, with the response amplitude serving as the predictor variable. The area under the ROC curve (AUC) was computed and transformed to the LMI using LMI = (AUC -0.5) × 2. To assess the statistical significance of each LMI, a permutation test was performed by shuffling the class labels (pre-versus post-learning) 1,000 times and recomputing the AUC for each shuffle. The p-value for each cell was defined as the percentile of the observed AUC within the shuffled distribution. Cells with p ≥ 0.975 were classified as having a significantly positive LMI (LMI+), while cells with p ≤ 0.025 were classified as having a significantly negative LMI (LMI-).

To compare the LMI distributions with chance (Figure 3–figure supplement 1C), we obtained a null distribution by shuffling the pre-and post-learning labels. With mice as the unit, we computed in each mouse the fraction of neurons with a significant LMI, of which 5% are expected by chance, and the fractions of LMI+ and LMI-neurons (2.5% expected each) and tested them against these values across mice.

To measure day-to-day drift (Figure 3–figure supplement 4), the LMI was computed in the same way between two consecutive days, with one day on each side: between the two baseline days (Days −2 and −1), the only consecutive days without whisker learning, and across the first learning session (Days −1 and 0; Day-0 passive trials are delivered after the learning session). For each mouse and pair of days, we computed the fractions of LMI+ and LMI-neurons and the net fraction (LMI+ minus LMI-).

#### Response amplitude and population analyses of passive responses

Population responses before and after learning (Figures 3C–E and 4B–D, G–H) were computed for each mouse as the mean response across cells (or of its wS2-or wM1-projecting neurons), averaged over the passive trials of Days −2 and −1 (pre) and of Days +1 and +2 (post). In all analyses of projection neurons (Figure 4 and Figure 4– figure supplement 1), mice with fewer than three neurons of a given type were excluded from the analysis of that type.

To test whether responses adapted within the passive block (Figure 3–figure supplement 3), the population response of each trial was averaged in bins of 10 trials along the last 40 passive trials of each day, and over the first 20 and the last 20 of these trials (trials 11–30 and 31–50 of the block). The comparison of pre-and post-learning responses was then repeated using only the first or only the last 20 trials of each day.

#### Similarity matrices and network reorganization index

Trial-by-trial similarity matrices were computed for each mouse using passive whisker trials. For each session (Days −2 to +2), the last 40 passive whisker trials were selected and the whisker-evoked response amplitude was averaged over 0-300 ms from stimulus onset for each trial. Cosine similarity was computed between all pairs of trials across the five imaging days, resulting in a 200 × 200 similarity matrix (5 days × 40 trials per day) for each mouse. Matrix entries were computed as the normalized dot product between trial pairs: for population activity vectors x and y, similarity = (x · y) / (||x|| × ||y||). A cosine similarity matrix was computed for each mouse and then averaged across mice. Within-day similarity was the mean similarity between the different trials of the same day.

To quantify the decorrelation between pre-and post-learning representations, a network reorganization index was defined for each mouse as the difference between the average within-period similarity and the average between-period similarity. Within-period similarity was calculated as the average similarity among all pre-learning trial pairs (Days −2 and −1) and the average similarity among all post-learning trial pairs (Days +1 and +2). Between-period similarity was calculated as the average similarity between all pre-learning and post-learning trial pairs. The network reorganization index was computed as: [((Within-pre similarity + Within-post similarity) / 2) -Between-pre-post similarity].

#### Signal power, noise power and response-pattern drift

Because cosine similarity between single trials depends on trial-to-trial variability, we decomposed each mouse responses on each day into signal and noise (Figure 3– figure supplement 5). For the N × T matrix of responses of N neurons in T = 40 trials, noise power was the trial-to-trial variance summed over neurons, and signal power the squared norm of the trial-averaged response vector (r̄) corrected for the contribution of noise, ||r̄||^2^ -noise power / T. For each mouse, signal and noise power were averaged over Days −2 and −1 (pre) and over Days +1 and +2 (post).

To compare response patterns across days independently of trial-to-trial noise (Figure 3–figure supplement 4D), we computed the Pearson correlation across neurons between the mean response vectors of two consecutive days, and divided it by the geometric mean of the reliabilities of the two mean vectors. The reliability of the mean vector of a day was its split-half correlation (mean over 100 random splits of the trials into two halves), with the Spearman–Brown correction 2r / (1 + r). A noise-corrected correlation of 1 indicates the same response pattern on both days. This was computed between the two baseline days (Days −2 and −1) and across the first learning session (Days −1 and 0), and, within each day, between the first and the last 20 of the 40 passive trials (each half’s reliability computed from its own 20 trials).

#### Decoding of pre-and post-learning and projection on the learning dimension

To test whether population activity patterns could discriminate pre-from post-learning representations, a logistic regression classifier was trained on passive whisker trials for each mouse. The whisker-evoked response amplitude (ΔF/F₀) was averaged over 0-300 ms from stimulus onset for each trial. Pre-learning trials (Days −2 and −1 pooled) were labelled as class 0 while post-learning trials (Days +1 and +2 pooled) were labelled as class 1. Features were z-scored across trials with the mean and standard deviation of the training set, and a logistic regression classifier with L2 regularization was trained using 10-fold stratified cross-validation. Decoding accuracy was computed as the mean accuracy across folds.

To evaluate whether Day 0 activity patterns resembled pre-or post-learning representations, the decoder was applied to all pairwise combinations of days (Days -2, −1, 0, +1, +2) using held-out test folds. For each mouse, the trials of each day were split into 10 folds. At each fold iteration, a decoder was trained exclusively on pre-learning (Days −2, −1) versus post-learning (Days +1, +2) trials from the training folds, then tested on all pairwise day combinations using the corresponding test folds. For each day pair (day_i_, day_j_), trials from day_i_ were assigned to class 0 and trials from day_j_ were assigned to class 1, and classification accuracy was computed. Accuracies were averaged across folds, and the pre-learning vs Day 0 and Day 0 vs post-learning accuracies of each mouse were the means over Days −2 and −1 and over Days +1 and +2, respectively. To test how much decoding relied on learning-modulated neurons (Figure 5–figure supplement 1B), the most modulated neurons (in steps of 5%) were removed before repeating the cross-validated decoding.

To track the progressive realignment of population representations during learning, whisker-evoked activity from Day 0 task trials was projected onto the learning axis defined by the pre-versus post-learning decoder. For each mouse, a fixed decoder was trained, as above, on all passive whisker trials of Days −2 and −1 (pooled) versus Days +1 and +2 (pooled), excluding Day 0, using activity averaged over 0–300 ms after stimulus onset. The sign of the decision function was set so that positive values indicate activity closer to post-learning responses and negative values activity closer to pre-learning responses, 0 being the decoder boundary. The decoder was then applied to all Day 0 whisker trials of the task. For these trials, the response was averaged over 0–180 ms from stimulus onset, which ends before the first lick on nearly all trials (first lick before 180 ms on 0.6% of whisker hits in R+ mice and 3.1% in R-mice), so that lick-related activity contributes minimally to the projection. Responses were z-scored with the training-set parameters and projected with the decision function; projections were averaged in a sliding window of 10 consecutive whisker trials. As reference values, the passive pre-and post-learning trials used to train the decoder were projected in the same way (Figure 5G): each trial was projected by a decoder trained on the other folds of a stratified 10-fold split (training on the 0-300 ms window), using its response averaged over 0-180 ms, as for the active Day 0 trials.

#### Detection of reactivations

To detect spontaneous reactivations of whisker-evoked activity patterns, a template-matching approach was applied to the correct-rejection catch trials of each session, i.e. catch trials (no auditory or whisker stimulus) without a lick in the response window. Each trial was analyzed from 1 s before to 6 s after trial onset; mice did not lick in the no-lick period preceding trial onset nor in the 1-s response window, but could lick after it. For each mouse and day, a whisker response template was first constructed from passive whisker trials. The template was defined as the trial-averaged whisker-evoked population activity vector, computed by averaging the response amplitude of each neuron over 0–300 ms from stimulus onset across the last 40 passive trials of that day.

Neural activity (ΔF/F_0_, without baseline subtraction) during correct-rejection trials was then compared to the template of the same day by computing, at each imaging frame, the Pearson correlation across neurons between the template and the population activity vector. Each trial was treated as a separate segment, since consecutive catch trials are not contiguous in time: the correlation time series of each trial was smoothed with a Savitzky–Golay filter (window length of 5 frames, polynomial order 2), and reactivation events were identified within each trial as peaks of the smoothed correlation exceeding a detection threshold, with a minimum inter-peak distance of 4 frames (133 ms) and a minimum prominence of 0.15 (scipy.signal.find_peaks).

To ensure that detected events were unlikely to arise from chance fluctuations, a detection threshold was determined for each mouse from surrogate data in which activity was shifted in time independently for each cell. For each of the two pre-learning days (Days −2 and −1), the concatenated correct-rejection activity of each neuron was circularly shifted by an independent random lag, the correlation time series with that day’s template was recomputed, and its 99^th^ percentile was taken; this was repeated 1,000 times per day. The detection threshold was the median of the 2,000 resulting percentiles, and was applied to all five days. The reactivation rate was the number of events divided by the total duration of the correct-rejection trials of the session (events per minute).

#### Participation in reactivations

To quantify each neuron’s engagement in reactivation events, a participation rate was computed for each neuron and day. A neuron was considered to participate in an event if its baseline-subtracted ΔF/F_0_, averaged over a window of ±4 frames (±133 ms) centered on the event, reached its participation threshold; events closer than 4 frames to a trial edge were not used. The participation threshold of each neuron was 2.5 times its noise SD, estimated on the same quantity: the ±4-frame mean ΔF/F_0_ at all time points of the correct-rejection trials at which events could be counted, pooled over the five days. The noise SD was estimated as 1.4826 x the median absolute deviation, a robust estimate that is insensitive to sparse calcium transients. The participation rate of a neuron on a day was the fraction of events from that day in which it participated.

To avoid circularity (a neuron contributes to the template correlation in proportion to its weight), participation was measured in held-out neurons. For each mouse, neurons were split at random into two halves. Reactivation events were detected using one half only (its template, its activity, and a detection threshold recomputed for that half from 500 circular-shift surrogates per pre-learning day, as above), and participation was measured in the other half. The halves were then swapped, and the procedure was repeated for 10 random splits, so that each neuron was held out 10 times and a neuron’s participation rate on a day was its mean over the splits. For comparison, participation was also measured, in the same events, in the half of the neurons that had detected them (in-sample).

To control for activity levels, we computed for each neuron and day a chance participation rate, the fraction of time points at which events could be counted at which its ±4-frame mean ΔF/F_0_ reached the threshold, i.e. the participation expected if events occurred at random times. Participation above chance, the participation rate minus the chance rate, was the main measure (Figures 6D–E); the participation rate itself is shown in Figure 6–figure supplement 2. Participation analyses included mice with at least three reactivation events on Day 0 (18 R+ and 14 R-mice; one R+ and two R-mice were excluded). Event-rate analyses (Figure 6C and its controls) included all mice. For Figure 6D, participation above chance was averaged per mouse over its LMI+ (or LMI-) neurons on each day, and its trend across days was quantified as the slope of a linear regression on day, one slope per mouse. For Figure 6E, Day 0 participation above chance was related to the LMI of each neuron with a linear mixed-effects model (see Statistics).

#### Controls for the reactivation analyses

To control for licking, reactivation rates were computed in 0.5-s bins across the trial and in two windows: the lick-free part of the trial (-0.6 to 1 s from trial onset) and the post-response part, in which licks were possible (1 to 5.6 s). The first and last 0.4 s of each trial were excluded from both windows to avoid edge artefacts. Each mouse’s rates were averaged over the five days (Figure 6–figure supplement 1A). In addition, the detection (including the surrogate thresholds) and the participation analyses were repeated using only the lick-free part of each trial (-1 to +1 s from trial onset) (Figure 6–figure supplement 3).

To test whether the detected events were specific to the whisker template, for each mouse and day, event detection was repeated with 1,000 templates in which the neuron identities of the real template were shuffled, using the same activity and the same detection threshold as for the real template. A specificity index was computed for each mouse and day as the real event rate minus the mean event rate of the shuffled templates (Figure 6–figure supplement 1B).

Finally, the analyses were repeated with detection thresholds at the 99.5^th^ and 99.9^th^ percentiles of the surrogates (events re-detected for all neurons and, in the held-out analysis, for each half and split), and with participation thresholds of 5 and 10 noise SD (Figure 6–figure supplement 3).

#### Statistics

All comparisons took mice as unit, with one value per mouse except where stated. We used non-parametric tests: the Mann–Whitney U test for comparisons between independent groups, the Wilcoxon signed-rank test for paired comparisons within mice and for comparisons of per-mouse values with a reference value (0, or the fraction expected by chance), and the Friedman test for changes across the five days. Trends across days or trials (Figure 5H, Figure 6D, Figure 5–figure supplement 2, and Figure 6–figure supplement 2B) were tested by computing one regression slope per mouse and comparing the slopes with 0 (Wilcoxon signed-rank test). Tests were two-sided, except the tests of the Day 0 projection slope and of its correlation with behavior against 0 (Figure 5H&I and Figure 5–figure supplement 2), which were one-sided.

Distributions of single-neuron values (LMI, decoder weights) were compared with the two-sample Kolmogorov–Smirnov distance between the pooled neurons of the two groups, with a p-value from a permutation test that shuffled whole mice between groups (R+ vs R−) or labels within mice (wS2p vs wM1p). The p-value was (1 + number of permutations with a distance at least equal to the observed one) / (1 + 10,000 permutations).

The relationship between Day 0 participation and LMI across neurons (Figure 6E, Figure 6–figure supplement 2A, Figure 6–figure supplement 3B) was tested with a linear mixed-effects model, participation ∼ LMI + (1 + LMI | mouse), with a random intercept and a random slope of LMI per mouse, fitted with statsmodels (MixedLM; the fit with the highest likelihood across several optimizers was kept). The fixed-effect slope is reported with its 95% confidence interval and a Wald test.

Trial-by-trial comparisons of learning curves (Figure 1D and Figure 1–figure supplement 1D&E) were corrected for multiple comparisons across trials with the Benjamini–Hochberg procedure. Other tests were performed per day or per condition and are reported without correction. Bars and shaded areas show the mean and the bootstrapped 95% confidence interval across mice (1,000 resamples) throughout.

## Data availability

Imaging data, behavioral data, and calcium traces were combined into a single NWB file for each behavioral session. NWB offers a common format for sharing and analyzing neurophysiology data (Rübel et al., 2022). The NWB files are available on the DANDI archive (https://dandiarchive.org/dandiset/002039). The metadata and preprocessed data needed to reproduce the figures will be available on Zenodo (https://doi.org/10.5281/zenodo.24107306).

## Code availability

Code for data acquisition and behavior control is available on Github (https://github.com/LSENS-BMI-EPFL/behavior_control). The code used to preprocess and convert the data into NWB format is available on Github (https://github.com/LSENS-BMI-EPFL/NWB_converter). The code used for data analysis, which generates all figure panels and statistics from the NWB files, is available on Github (https://github.com/LSENS-BMI-EPFL/renard_foustoukos) and will be available on Zenodo (https://doi.org/10.5281/zenodo.24107306).

## Acknowledgements

We thank James Priestley and the members of the Petersen laboratory for helpful discussions. This work was supported by grants 310030_219343 (S.C. and C.C.H.P.) and TMAG-3_209271 (C.C.H.P.) from the Swiss National Science Foundation.

## Author contributions

A.R., G.F., S.C. and C.C.H.P. conceptualized the study; A.R., G.F., R.F.D., A.B. and P.B. wrote the data acquisition codes. A.R., G.F. and M.I. performed all animal experiments and constructed the database. A.R. and G.F. analyzed the data. A.R., S.C. and C.C.H.P. wrote and edited the manuscript, with comments from all authors; S.C. and C.C.H.P. acquired funding and provided overall supervision.

---

**Figure 1–figure supplement 1.**
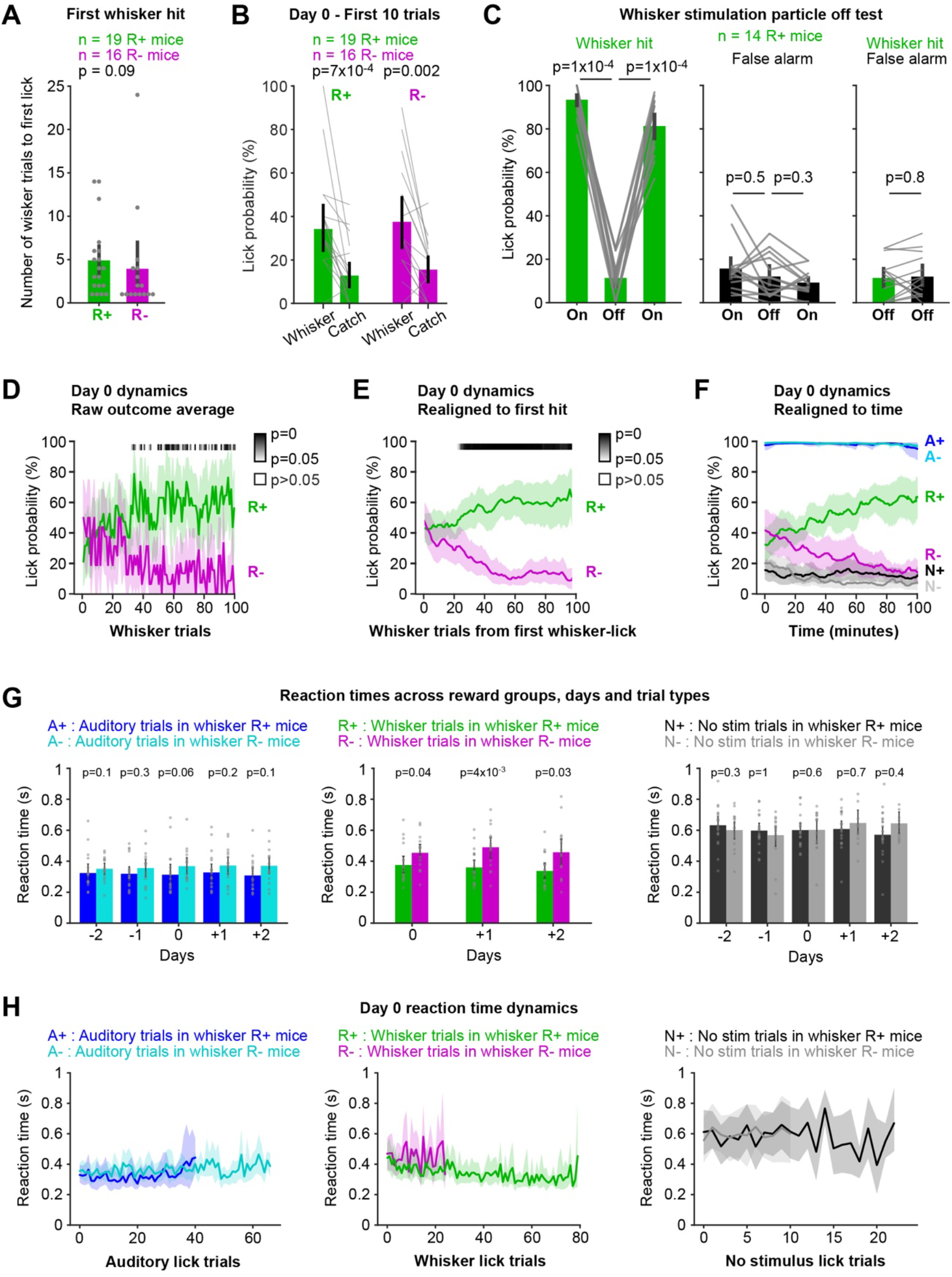
Behavioral quantification of rapid whisker learning. (**A**) Whisker trial number of the first whisker hit on Day 0 (1, first whisker trial), R+ (n = 19 mice) vs R-mice (n = 16 mice). Bars, mean ± 95% CI; dots, mice; p-value, Mann-Whitney U test. (**B**) Lick probability in the first 10 whisker trials of Day 0, and in the catch trials delivered before the 10th whisker trial. Bars, mean ± 95% CI; lines, mice; p-values, Wilcoxon signed-rank test, whisker vs catch trials, within each group (n = 19 R+ and 16 R-mice). Whisker lick probability did not differ between R+ and R-mice (Mann-Whitney U test). (**C**) Particle-off control on Day +3 in 14 R+ mice. The metal particle was removed from the C2 whisker to control for magnetic stimulus artefacts. Lick probability is shown in a first particle-on block (On), a particle-off block (Off) and a second particle-on block (On). Left: whisker hit rate. Middle: catch-trial false-alarm rate. Right: whisker hit rate and false-alarm rate in the particle-off block. Bars, mean ± 95% CI; grey lines, mice; p-values, Wilcoxon signed-rank test (n = 14 mice). (**D**) Whisker lick probability across Day 0 whisker trials, averaged from the binary single-trial outcomes, for R+ (green) and R-(magenta) mice. Lines, mean ± 95% CI across mice (n = 19 R+ and 16 R-mice). The grey scale bar gives the p-value of a Mann-Whitney U test, R+ vs R-, at each whisker trial, corrected across trials (Benjamini-Hochberg). (**E**) As in panel D, but with lick probability estimated by the state space model (as in Figure 1D) and whisker trials realigned to each mouse’s first whisker lick. (**F**) Lick probability on Auditory (A+, dark blue in R+ mice; A-, light blue in R-mice), whisker (R+, green in R+ mice; R-, magenta in R-mice) and no-stimulus catch trials (N+, black in R+ mice; N-, grey in R-mice) as a function of time on Day 0 (minutes from session start). Lines, mean ± 95% CI across mice. (**G**) Mean reaction time on lick trials across days, for auditory (left), whisker (middle) and catch trials (right; i.e. false alarms) in R+ and R-mice. Reaction time was measured from stimulus onset to the first lick. Bars, mean ± 95% CI; dots, mice; p-values, Mann-Whitney U test, R+ vs R-, on each day (n = 19 R+ and 16 R-mice, except mice without lick trials of that type on that day: 15 R-mice for whisker trials on Day +2, and 14, 13 and 11 R-mice for catch trials on Days 0, +1 and +2). (**H**) Mean reaction time across the successive lick trials of Day 0 for each trial type. Lines, mean ± 95% CI across mice.

**Figure 2–figure supplement 1.**
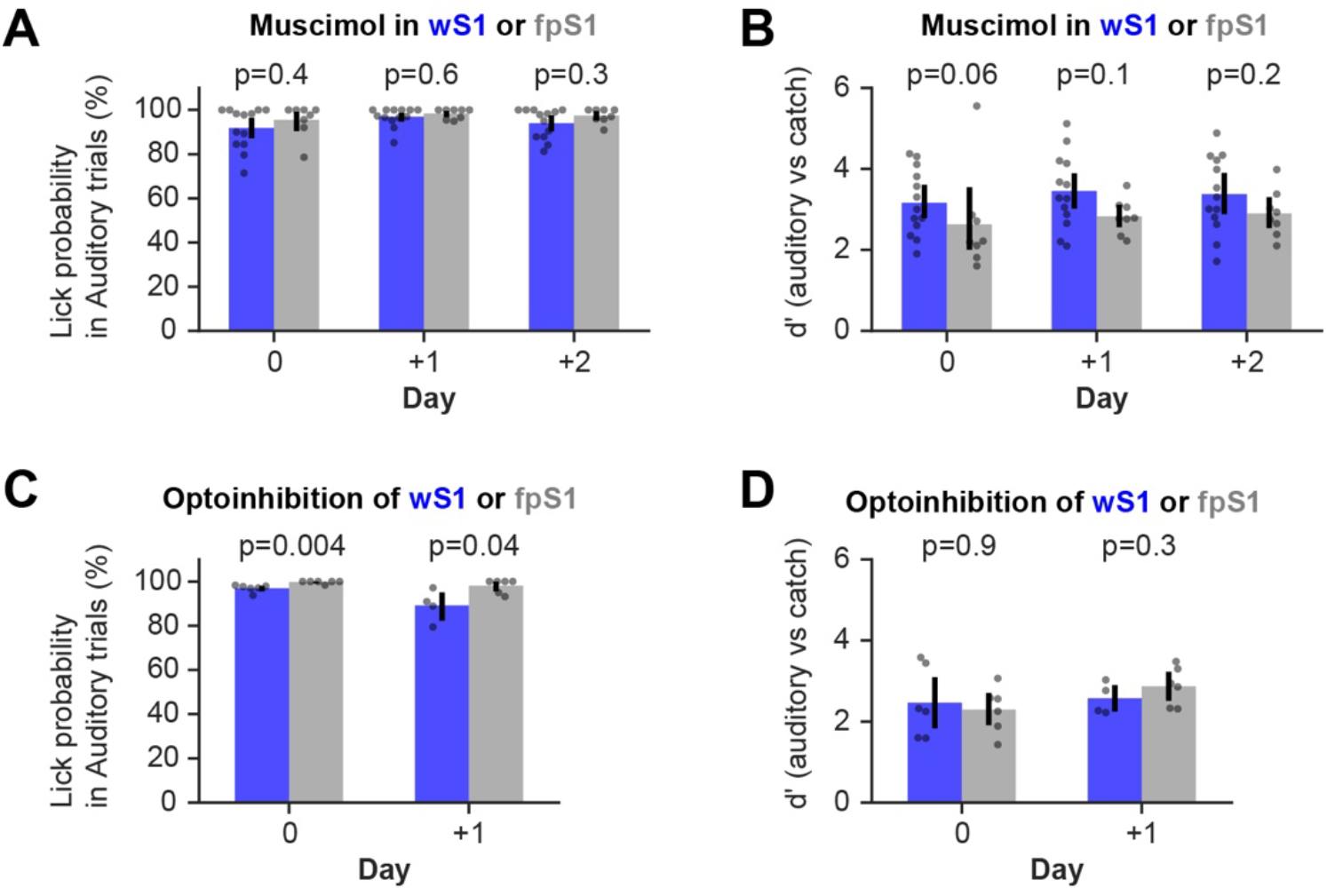
Auditory performance during inactivation of the barrel cortex during whisker learning. (**A**) Auditory lick probability of the mice of Figure 2, for inactivation of wS1 (blue) and fpS1 (grey) using muscimol on Days 0, +1 and +2 (n = 13 wS1 and 8 fpS1 mice). (**B**) Same as panel A, but for auditory d′ (auditory vs catch trials). (**C**) Auditory lick probability of the mice during optogenetic inactivation of wS1 (blue) and fpS1 (grey), on Day 0 (light) and Day +1 (no light) (n = 6 wS1 and 6 fpS1 mice; but only 4 wS1 mice on Day +1). (**D**) Same as panel C, but for auditory d′ (auditory vs catch trials). In all panels: bars, mean ± 95% CI; dots, mice; p-values, Mann-Whitney U test, wS1 vs fpS1, on each day.

**Figure 2–figure supplement 2.**
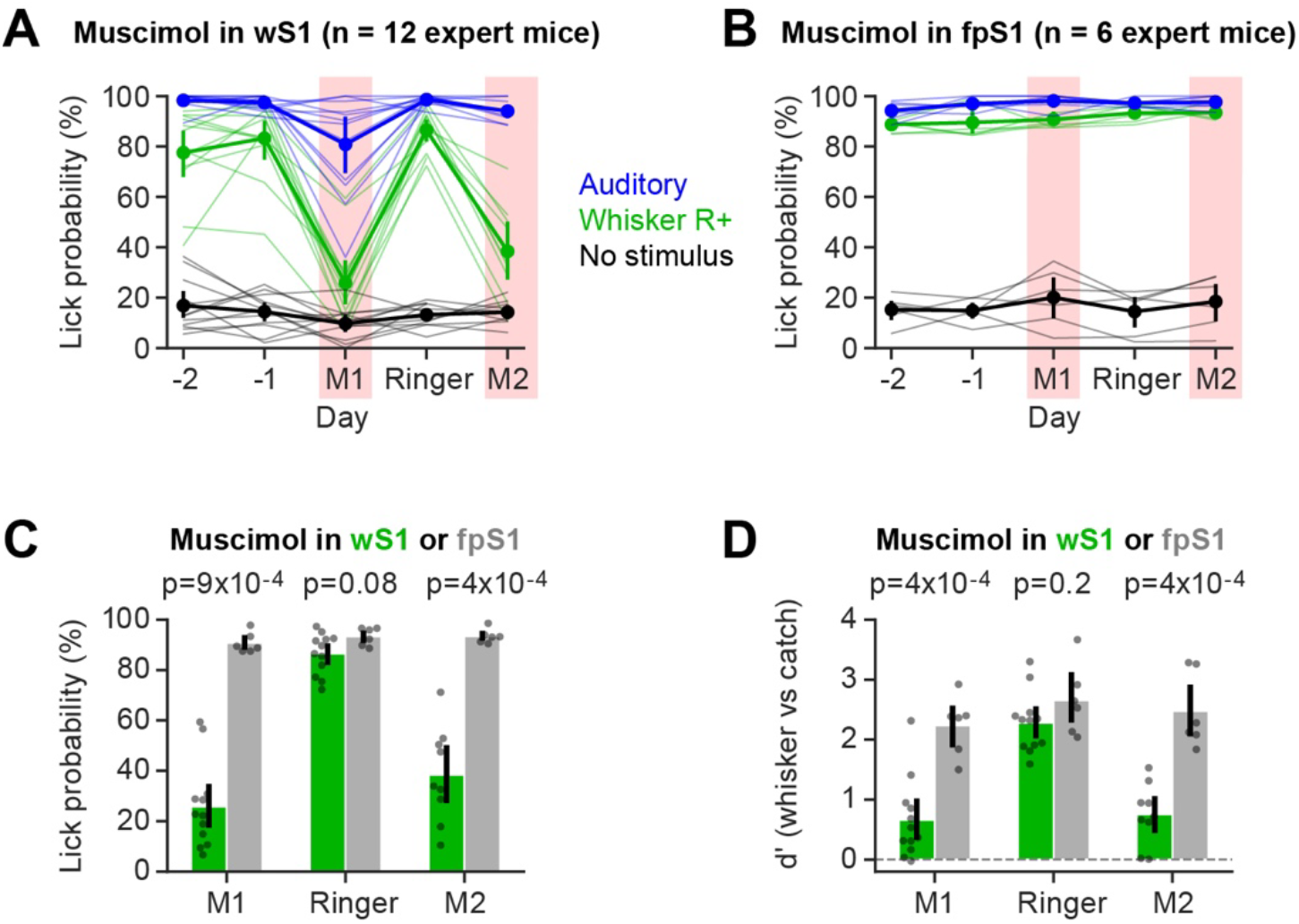
Barrel cortex inactivation also impairs whisker-guided licking in expert mice. (**A**) A separate cohort of R+ mice was trained to expert performance in the whisker task. After two high-performance baseline days (Day −2 and −1), muscimol was injected in wS1 on a first day (M1), Ringer’s solution on the next day, and muscimol again on a third day (M2). Lick probability on auditory (blue), whisker (green) and catch trials (black) across days for wS1 inactivation (n = 12 expert mice, except Day M2 where n = 9 expert mice). Muscimol days are shaded in red; thin lines, mice; points, mean ± 95% CI. (**B**) Same as panel A, but for fpS1 inactivation (n = 6 expert mice). (**C**) Whisker lick probability for inactivation of wS1 (green) vs fpS1 (grey) on the muscimol and Ringer days. Bars, mean ± 95% CI; dots, mice; p-values, Mann-Whitney U test, wS1 vs fpS1, on each day (n = 12 wS1 and n = 6 fpS1 expert mice; except Day M2 where n = 9 wS1 expert mice). (**D**) Same as panel C, but for d′ (whisker vs catch trials).

**Figure 2–figure supplement 3.**
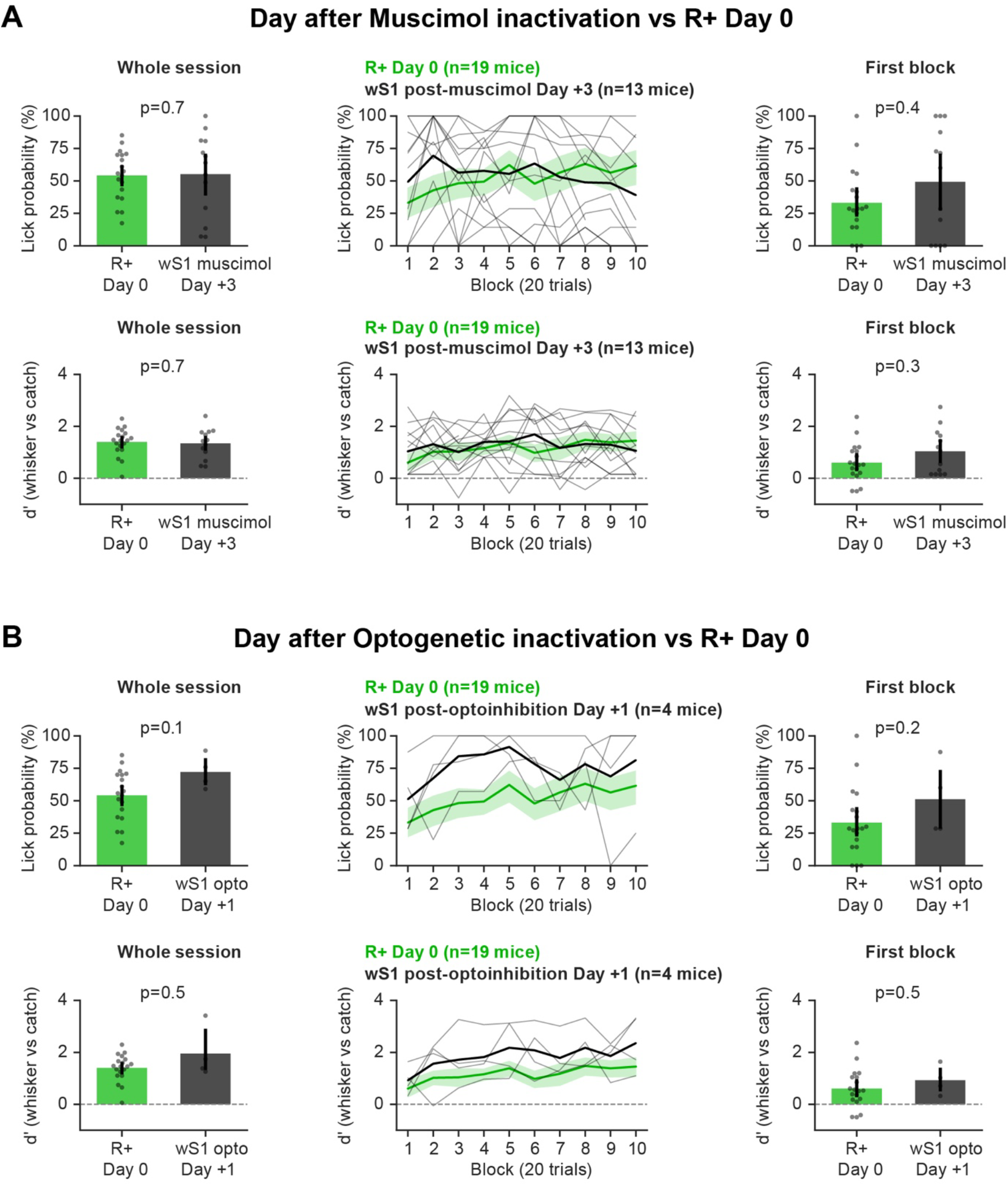
Performance on the control day after inactivation compared with naive mice learning on Day 0. (**A**) wS1 muscimol mice on Day +3, their first day without muscimol (n = 13 mice, black), compared with the naive R+ imaging mice on Day 0 (n = 19 mice, green). Top: whisker lick probability. Bottom: d′ (whisker vs catch trials). Left: whole session. Middle: first ten blocks of 20 trials. Naive mice are shown as mean ± 95% CI; inactivated mice are shown individually (thin lines) and as their mean (thick line). Right: first block of 20 trials. Bars, mean ± 95% CI; dots, mice; p-values, Mann-Whitney U test, inactivated vs naive mice. (**B**) As in panel A, but for wS1 optogenetic inactivation mice on Day +1, their first day without light after the inactivation Day 0 (n = 4 mice; 2 of the 6 wS1 mice were not trained on Day +1).

**Figure 3–figure supplement 1.**
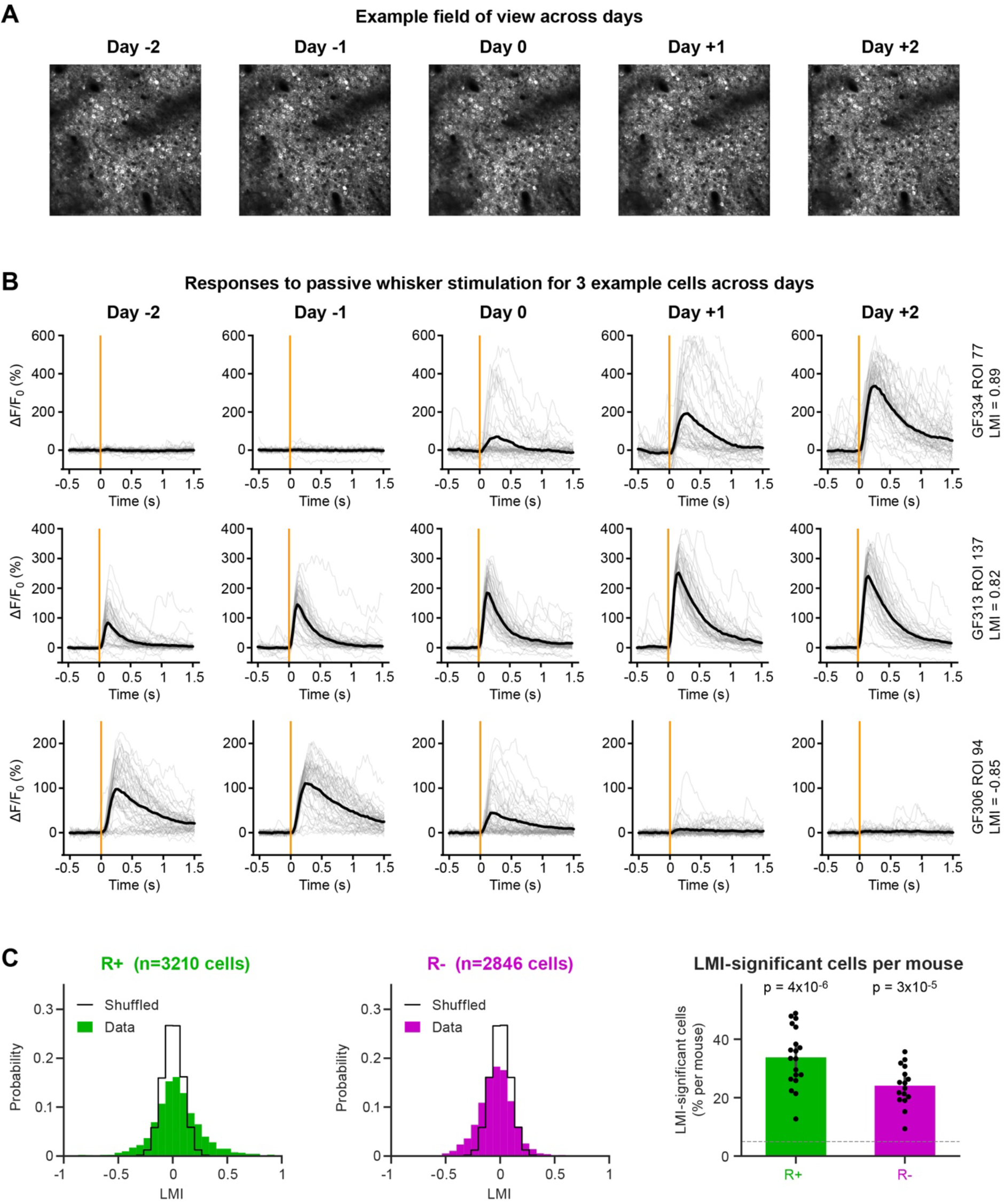
Imaging stability, example learning-modulated neurons and comparison of the LMI with chance. (**A**) Field of view of an example mouse on the five imaging days (mean of the first 1,000 frames of each day after motion correction). The recordings of the five days were registered together and each region of interest was defined once for all days (see Methods). (**B**) Responses to the passive whisker stimulus on the five days, for three example neurons from three R+ mice. Top: a neuron with a strong positive LMI that did not respond before learning. Middle: a neuron with a strong positive LMI whose response was enhanced. Bottom: a neuron with a strong negative LMI whose response was abolished. Grey, single trials; black, mean; orange line, stimulus onset. (**C**) Left and middle: distribution of the LMI of all neurons (filled; R+, n = 3,210 neurons from 19 mice; R-, n = 2,846 neurons from 16 mice) and null distribution obtained by shuffling the pre-and post-learning labels (black outline; LMI of every neuron in every shuffle pooled). Right: percentage of neurons with a significant LMI in each mouse. Dashed line, 5% expected by chance. Bars, mean ± 95% CI; dots, mice; p-values, Wilcoxon signed-rank test against 5% (n = 19 R+ and 16 R-mice).

**Figure 3–figure supplement 2.**
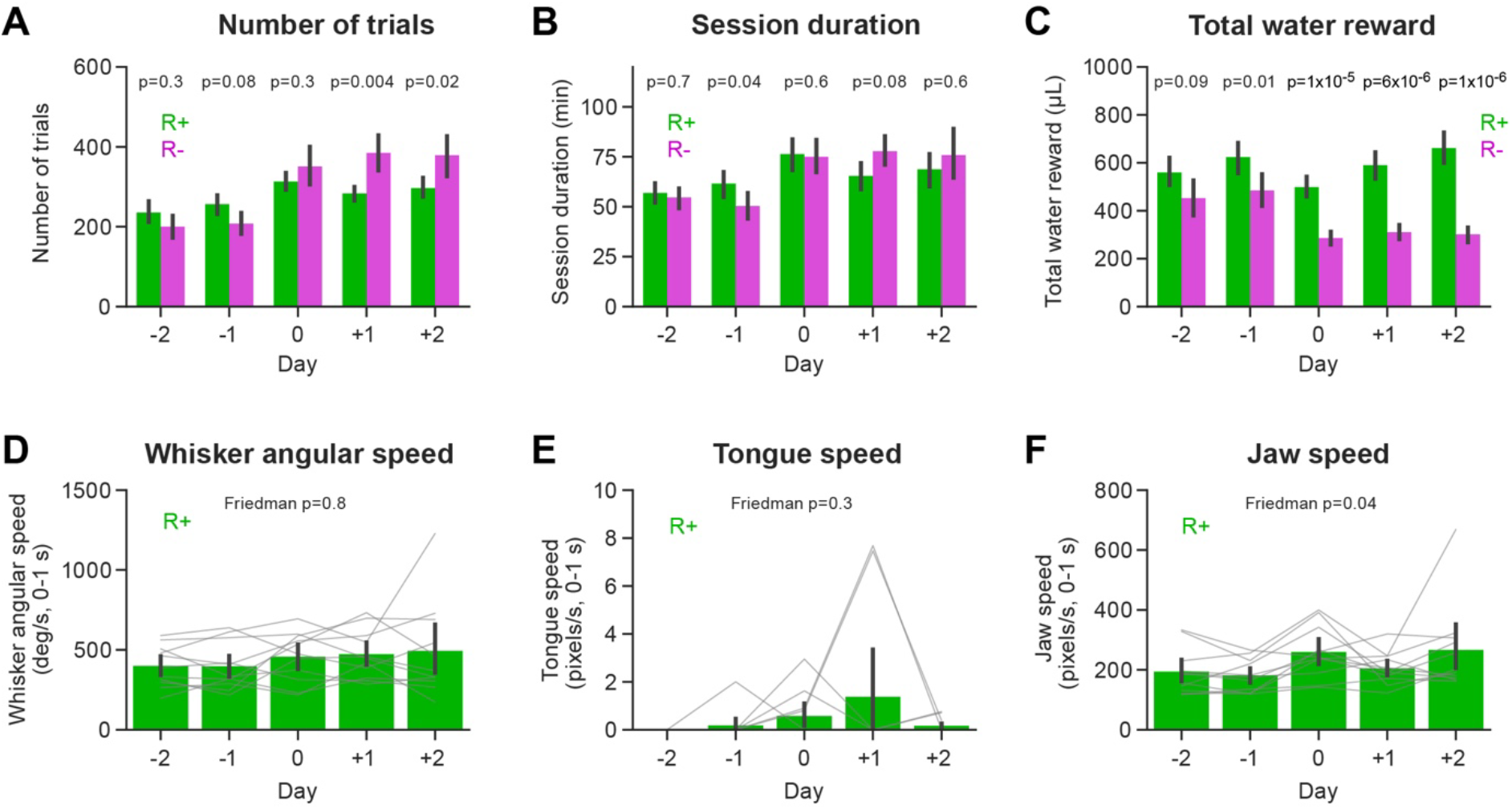
Behavioral state and orofacial movements of the imaged mice across days. (**A**) Number of trials in each active session across days for R+ mice (n = 19 mice, green) and R-mice (n = 16 mice, magenta). Bars, mean ± 95% CI; p-values, Mann-Whitney U test, R+ vs R-on each day. (**B**) Same as panel A, but for session duration (time from the first to the last task trial in minutes). (**C**) Same as panel A, but for total water reward (5 µL per rewarded trial). (**D**) Whisker angular speed in the 1 sec after passive whisker stimulus onset, averaged over the last 40 passive trials of each day, in the R+ mice with video recordings (n = 11 mice). Movements were tracked with DeepLabCut. Bars, mean ± 95% CI; grey lines, mice; p-value, Friedman test across days. (**E**) Same as panel D, but for tongue speed. (**F**) Same as panel D, but for jaw speed.

**Figure 3–figure supplement 3.**
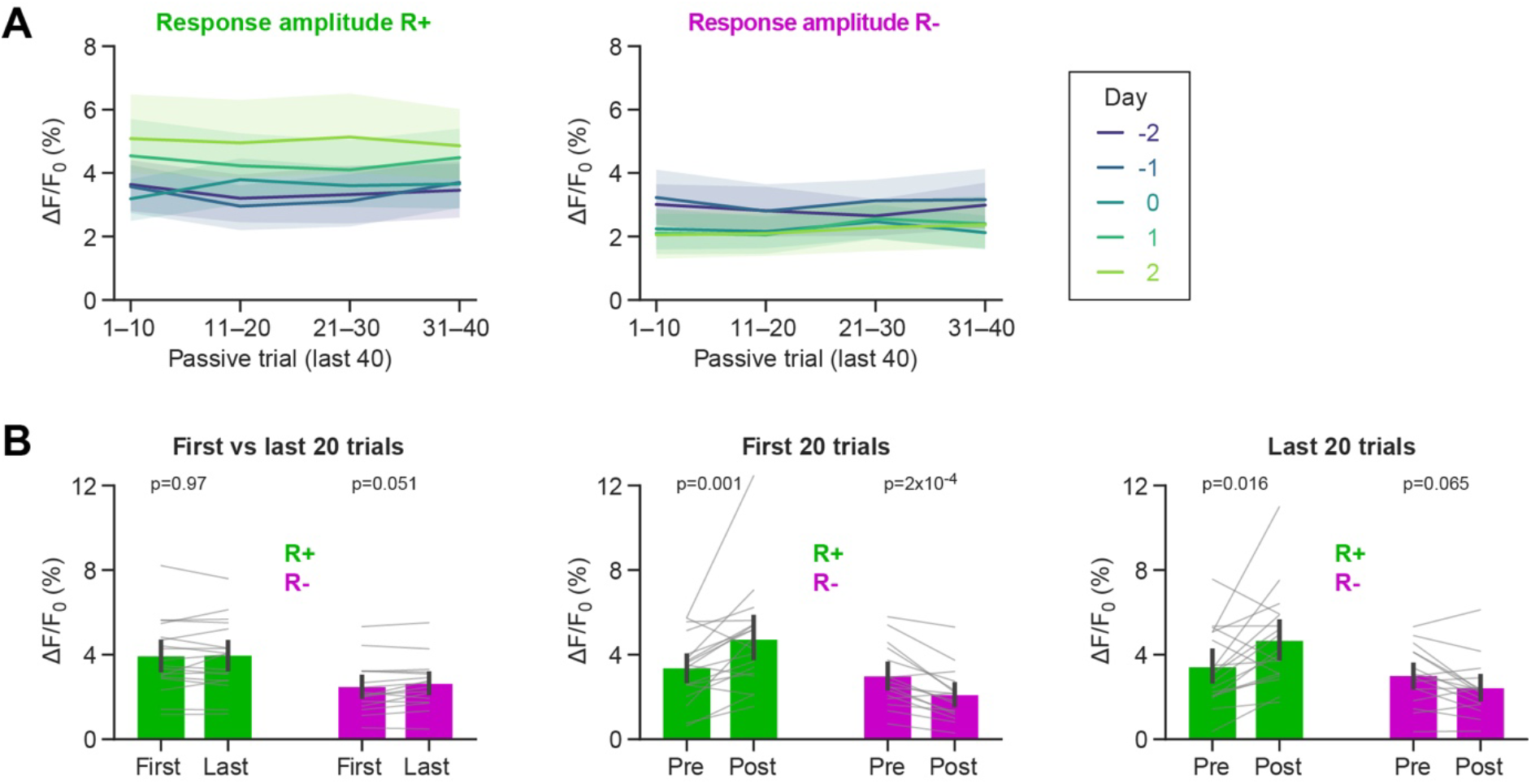
Stability of the passive whisker responses within the passive block. (**A**) Responses are from the last 40 of the 50 passive trials of each day, in block order, as the mean ΔF/F_0_ of each neuron over 0-300 ms after stimulus onset. Population response (mean over neurons per mouse) in bins of 10 trials along the passive block, on each day, for R+ (left) and R-mice (right) (n = 18 R+ and 16 R-mice). Lines, mean ± 95% CI across mice. (**B**) Left: population response in the first and the last 20 trials, each mouse averaged over the five days. Middle: population response before (Days −2 and −1; Pre) and after learning (Days +1 and +2; Post), computed from the first 20 trials of each day. Right: population response before (Days −2 and −1; Pre) and after learning (Days +1 and +2; Post), computed from the last 20 trials of each day. Bars, mean ± 95% CI; grey lines, mice; p-values, Wilcoxon signed-rank test, n = 18 R+ and 16 R-mice.

**Figure 3–figure supplement 4.**
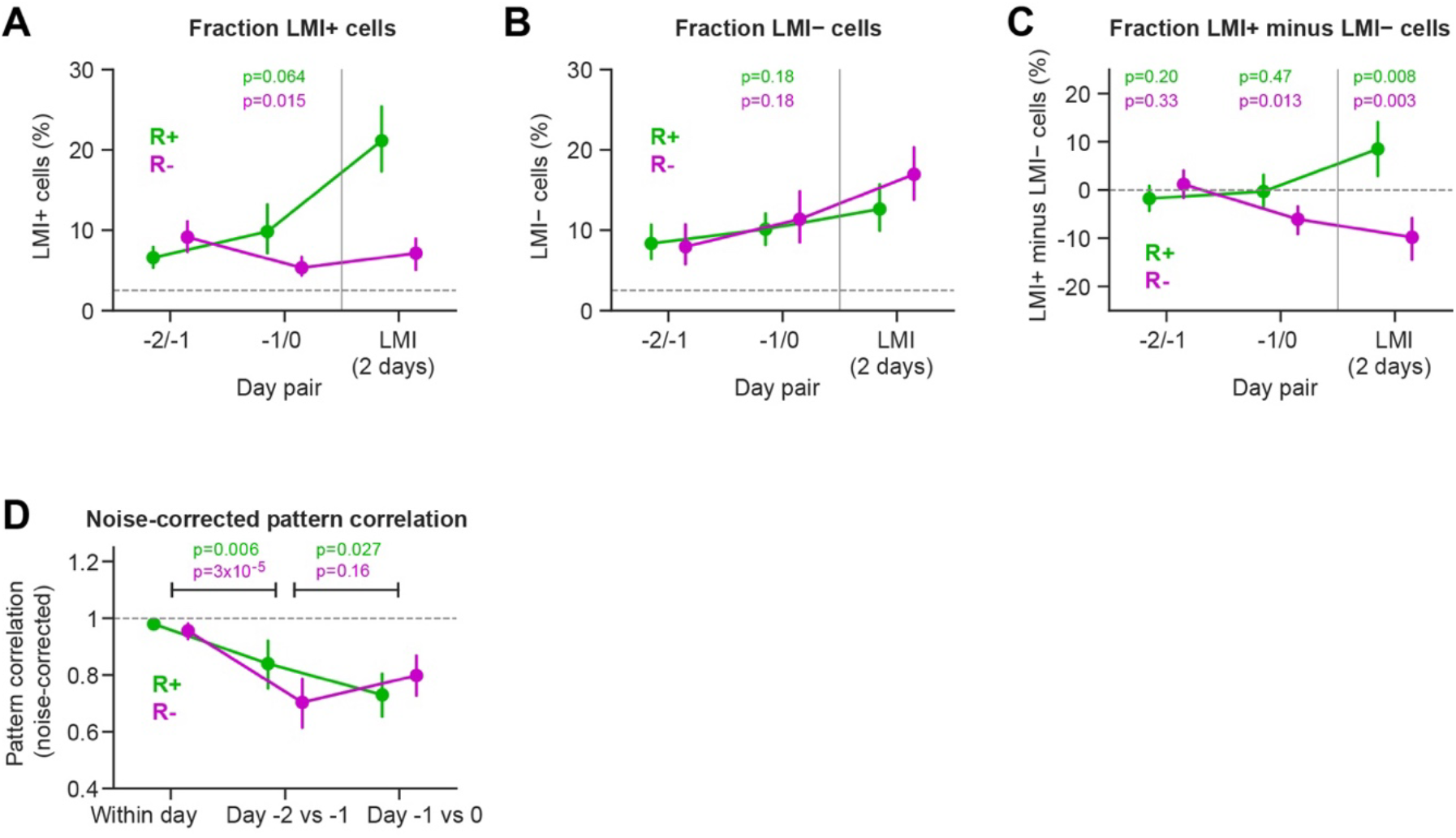
Reward learning-related plasticity *vs* day-to-day drift. (**A**) The percentage of LMI+ neurons. The LMI was computed between consecutive days, with one day on each side and the same shuffle test: between the two baseline days (-2/-1) and across the first learning session (-1/0) (n = 19 R+ and 16 R-mice). The LMI of Figure 3 (Days −2 and −1 vs Days +1 and +2) is shown for reference. Data points show mean ± 95% CI across mice. Dashed line, 2.5% expected by chance. P-value above Day pair −1/0 shows Wilcoxon signed-rank test of the change from −2/-1 to −1/0. (**B**) Same as panel A, but for LMI-neurons. (**C**) The percentage difference in LMI+ minus LMI-neurons. Data points show mean ± 95% CI across mice. Dashed line, 0% expected by chance. P-value above Day pair −1/0 shows Wilcoxon signed-rank test of the change from −2/-1 to −1/0. P-values above Day pair −2/-1 and the two-day LMI show Wilcoxon signed-rank test against 0. (**D**) Noise-corrected pattern correlation (correlation of the mean response vectors divided by their split-half reliabilities; 1 = identical pattern; see Methods). This is computed within the passive block (first vs last 20 trials; median over days), between Days −2 and −1, and between Days −1 and 0. Points, mean ± 95% CI; left p-values, Wilcoxon signed-rank tests within the block vs Days −2/-1; right p-values, Wilcoxon signed-rank tests Days −1/0 vs Days −2/-1.

**Figure 3–figure supplement 5.**
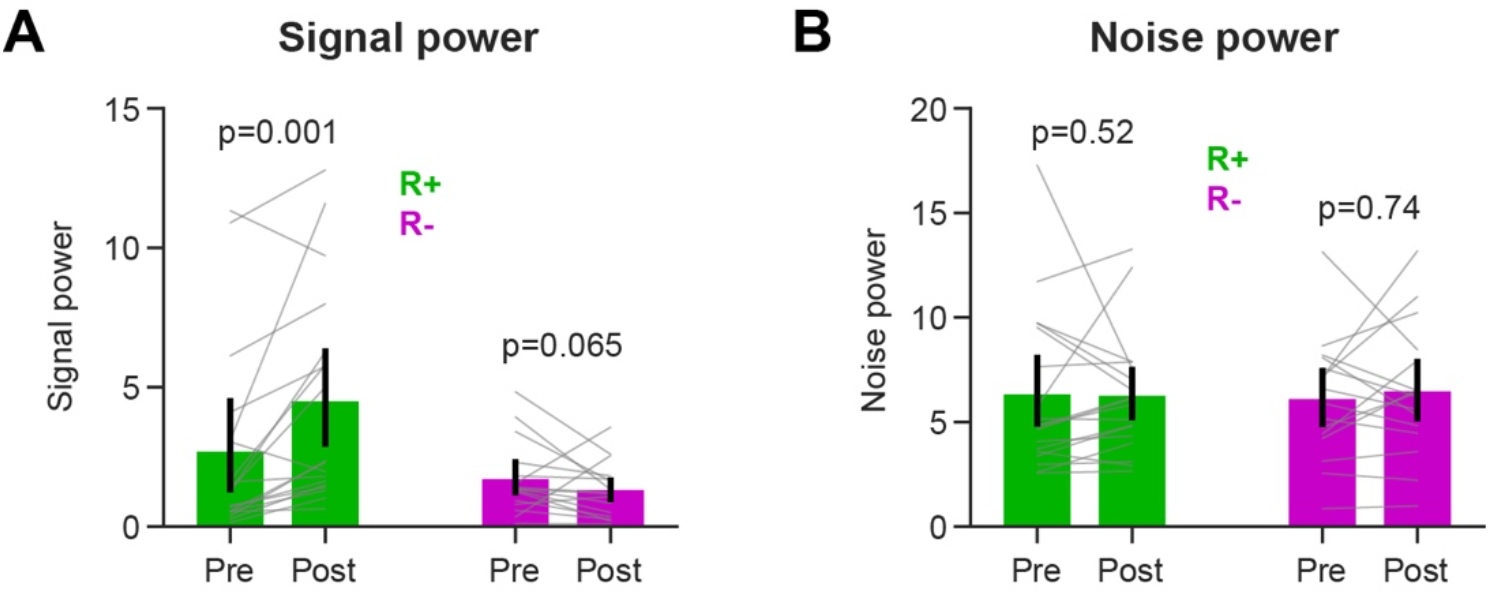
Signal and noise power of the passive whisker responses before and after learning. (**A**) Signal power: squared norm of the trial-averaged population response, corrected for the contribution of trial-to-trial noise. (**B**) Noise power: trial-to-trial variance of the responses, summed over neurons. Responses are from the last 40 passive trials of each day, as the mean ΔF/F_0_ of each neuron over 0-300 ms after stimulus onset. Values are averaged over Days −2 and −1 (Pre) and Days +1 and +2 (Post), for R+ (green; n = 18 mice) and R-mice (magenta; n = 16 mice). Bars, mean ± 95% CI; grey lines, mice; p-values, Wilcoxon signed-rank test, Pre vs Post.

**Figure 4–figure supplement 1.**
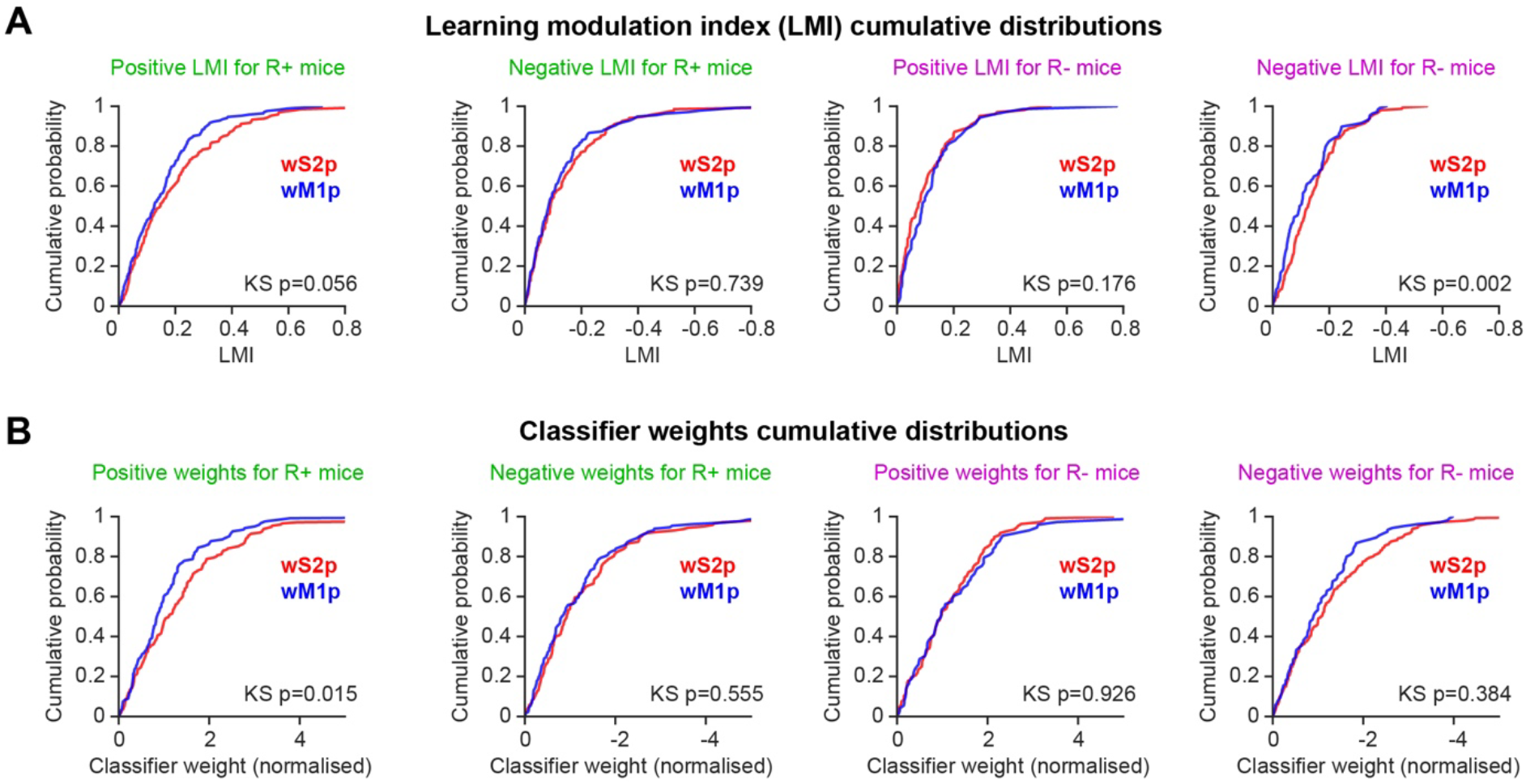
Comparison of wS2-and wM1-projecting neurons. (**A**) Cumulative distributions of the LMI of wS2p (red) and wM1p neurons (blue), shown separately for neurons with a positive and with a negative LMI, in R+ and R-mice. R+, positive LMI: 193 wS2p and 168 wM1p neurons; negative LMI: 134 and 135 neurons (18 mice). R-, positive LMI: 101 and 70 neurons; negative LMI: 152 and 108 neurons (12 mice). (**B**) As in panel A, but for the weights of the neurons in the pre-vs post-learning decoder (Figure 5). Weights are normalized by each mouse’s median absolute weight, because each mouse has its own decoder. R+, positive weights: 175 wS2p and 122 wM1p neurons; negative weights: 143 and 131 neurons (17 mice). R-, positive weights: 108 and 73 neurons; negative weights: 145 and 105 neurons (12 mice). The KS p-values indicate the Kolmogorov-Smirnov distance with a permutation test where the wS2p and wM1p labels were shuffled among the neurons of each mouse (10,000 permutations). As in Figure 4, only mice with at least three neurons of a type were included for that type.

**Figure 5–figure supplement 1.**
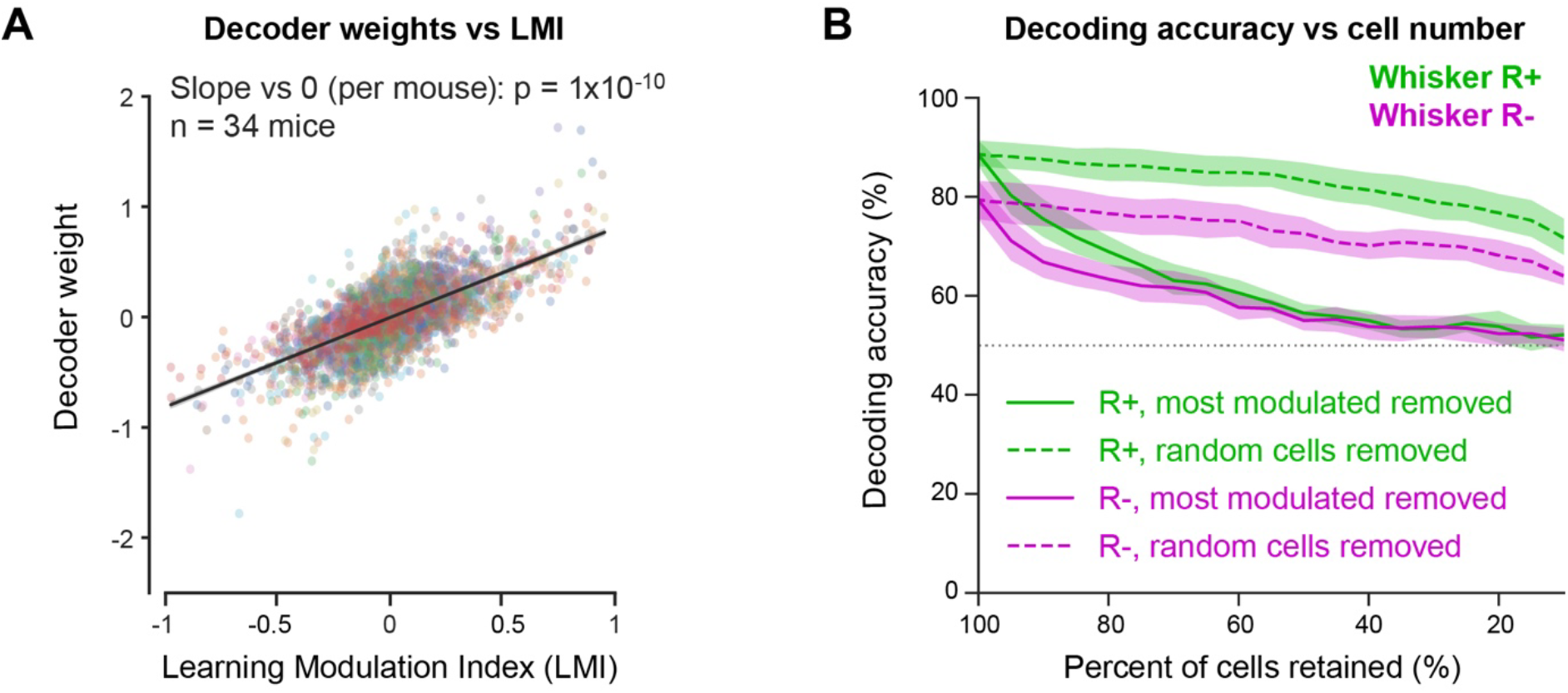
The decoder relies on learning-modulated neurons. (**A**) Weight of each neuron in the pre-vs post-learning decoder as a function of its LMI (n = 5,813 neurons from 34 mice; colors, mice). Line, linear regression on all neurons. Statistics: one regression slope per mouse, Wilcoxon signed-rank test against 0 (n = 34 mice). (**B**) Accuracy of the pre-vs post-learning decoder (10-fold stratified cross-validation) as a function of the percentage of neurons retained. Solid lines: within each fold, neurons were ranked by the absolute LMI computed on the training trials, and the most modulated ones were removed in steps of 5%. Dashed lines: removal of the same number of randomly chosen neurons. Lines, mean ± 95% CI across R+ mice (n = 18 mice) and R-mice (n = 16 mice). Dotted line, chance level (50%).

**Figure 5–figure supplement 2.**
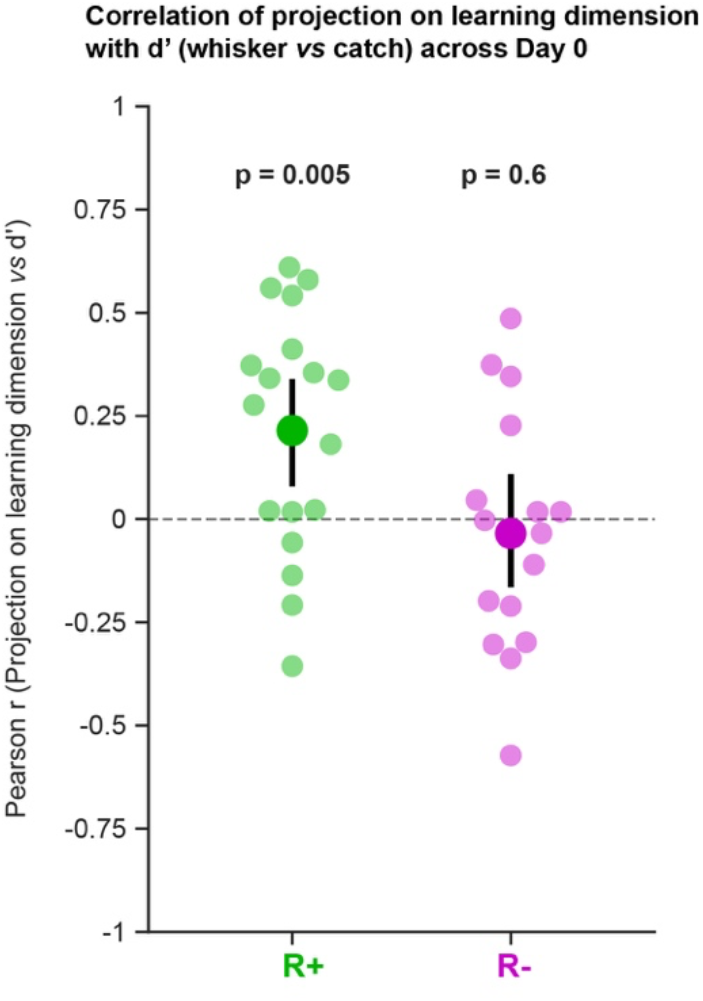
Correlation of the projection on the learning dimension with d′ across Day 0. Pearson correlation between the projection on the learning dimension (as in Figure 5I) and d′ (whisker vs catch trials) across Day 0 whisker trials, for each mouse. d′ was computed trial-by-trial from the state space estimates of the whisker and catch lick probabilities (as in Figure 1D). Small dots, mice; large dots, mean ± 95% CI; p-values, one-sided Wilcoxon signed-rank test against 0 (n = 18 R+ and 16 R-mice).

**Figure 6–figure supplement 1.**
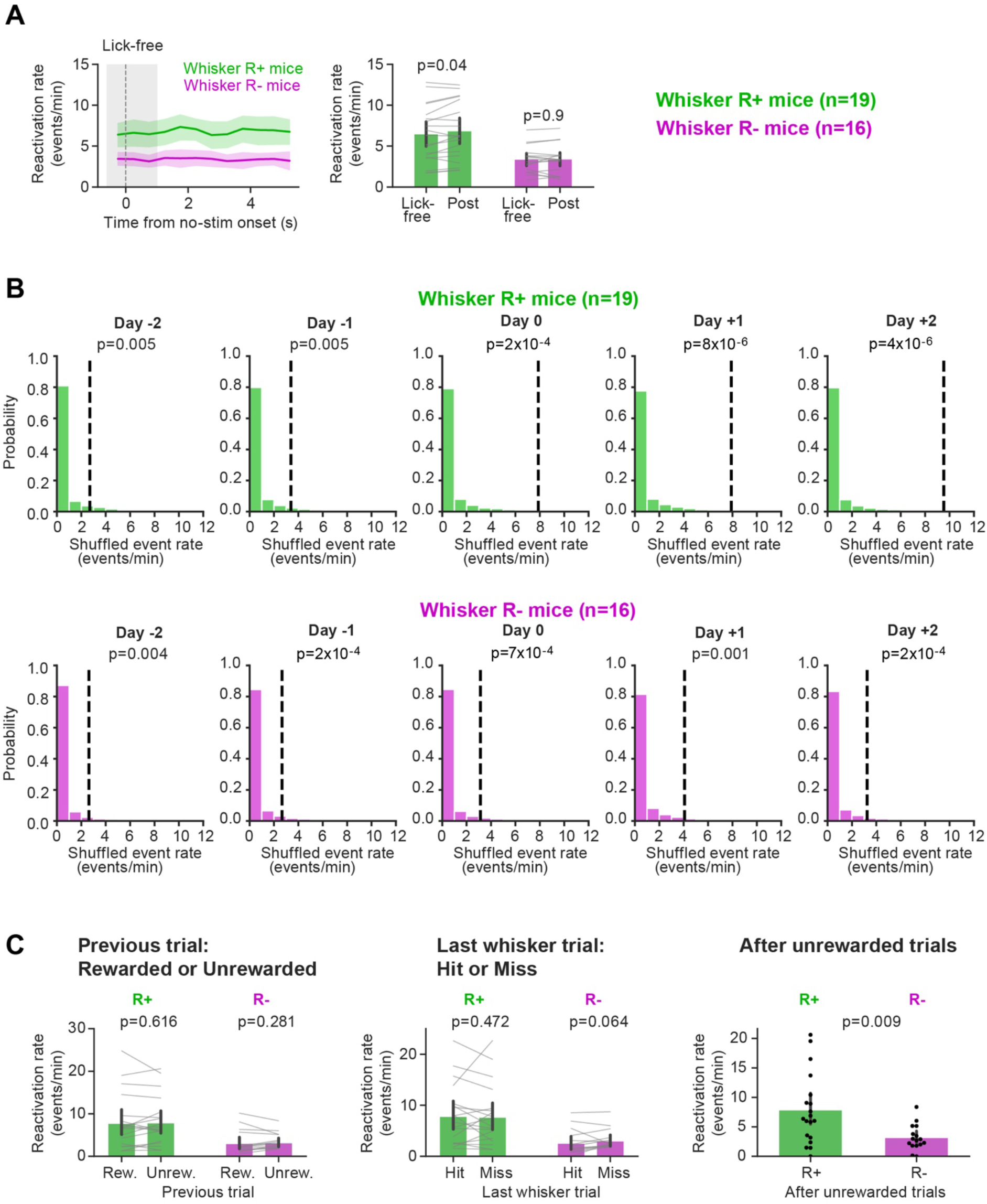
Controls for licking, template specificity, and previous trial outcome. (**A**) Left: Reactivation rate across the correct-rejection trials in 0.5-s bins from trial onset (no stimulus onset) for R+ mice (green; n = 19 mice) and R-mice (magenta; n = 16 mice) with each mouse averaged over the five days. Grey shading: lick-free epoch (no licks before trial onset or in the 1-s response window). The first and last 0.4 s of each trial are excluded, because events cannot be detected at the trial edges. Lines, mean ± 95% CI across mice. Right: Reactivation rate in the lick-free window (-0.6 to 1 s) and in the post-response window, in which mice could lick (1 to 5.6 s). Bars, mean ± 95% CI; grey lines, mice; p-values, Wilcoxon signed-rank test. (**B**) Event rates detected with 1,000 templates in which the identities of the neurons were shuffled, with the same activity and the same detection threshold as for the real template, on each day for R+ mice (above in green; n = 19 mice) and R-mice (below in magenta; n = 16 mice). Histograms, rates of all shuffled templates of all mice. Dashed lines, mean rate across mice with the real template. Statistics: for each mouse, the real rate minus the mean shuffled rate, Wilcoxon signed-rank test against 0 computed across mice. (**C**) Left: Day 0 reactivation rate of correct-rejection trials in R+ vs R-mice classified by the outcome of the previous trial (rewarded or unrewarded) (p-values, Wilcoxon signed-rank test within mice, n = 19 R+ and 16 R-mice). Middle: Day 0 reactivation rate of correct-rejection trials in R+ vs R-mice classified by the outcome of the last whisker trial (hit or miss) (p-values, Wilcoxon signed-rank test within mice, n = 19 R+ and 15 R-mice). Right: Day 0 reactivation rate of correct-rejection trials in R+ vs R-mice after unrewarded trials (p-value, Mann-Whitney U test, R+ vs R-). Categories with fewer than three trials in a mouse were not used. Bars, mean ± 95% CI; grey lines and dots, mice.

**Figure 6–figure supplement 2.**
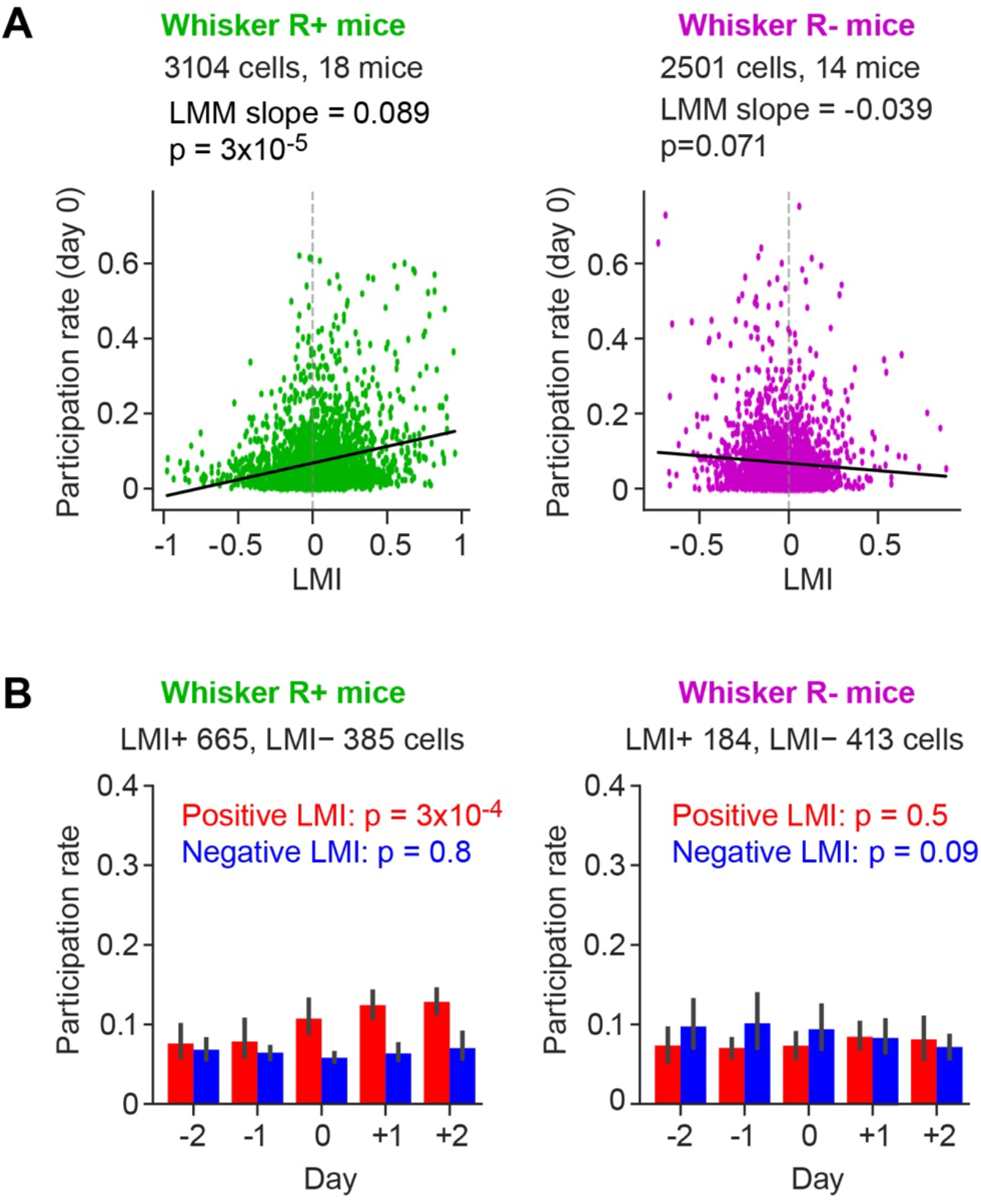
Participation rate in reactivations without correction for chance. (**A**) As in Figure 6E, but for the Day 0 participation rate of held-out neurons without correction for chance (R+, n = 3,104 neurons from 18 mice; R-, n = 2,501 neurons from 14 mice). Line, fixed effect of a linear mixed-effects model with a random intercept and a random slope per mouse. (**B**) As in Figure 6D, but for the participation rate without correction for chance. Bars, mean ± 95% CI across mice; one slope across days per mouse; p-values, Wilcoxon signed-rank test against 0.

**Figure 6–figure supplement 3.**
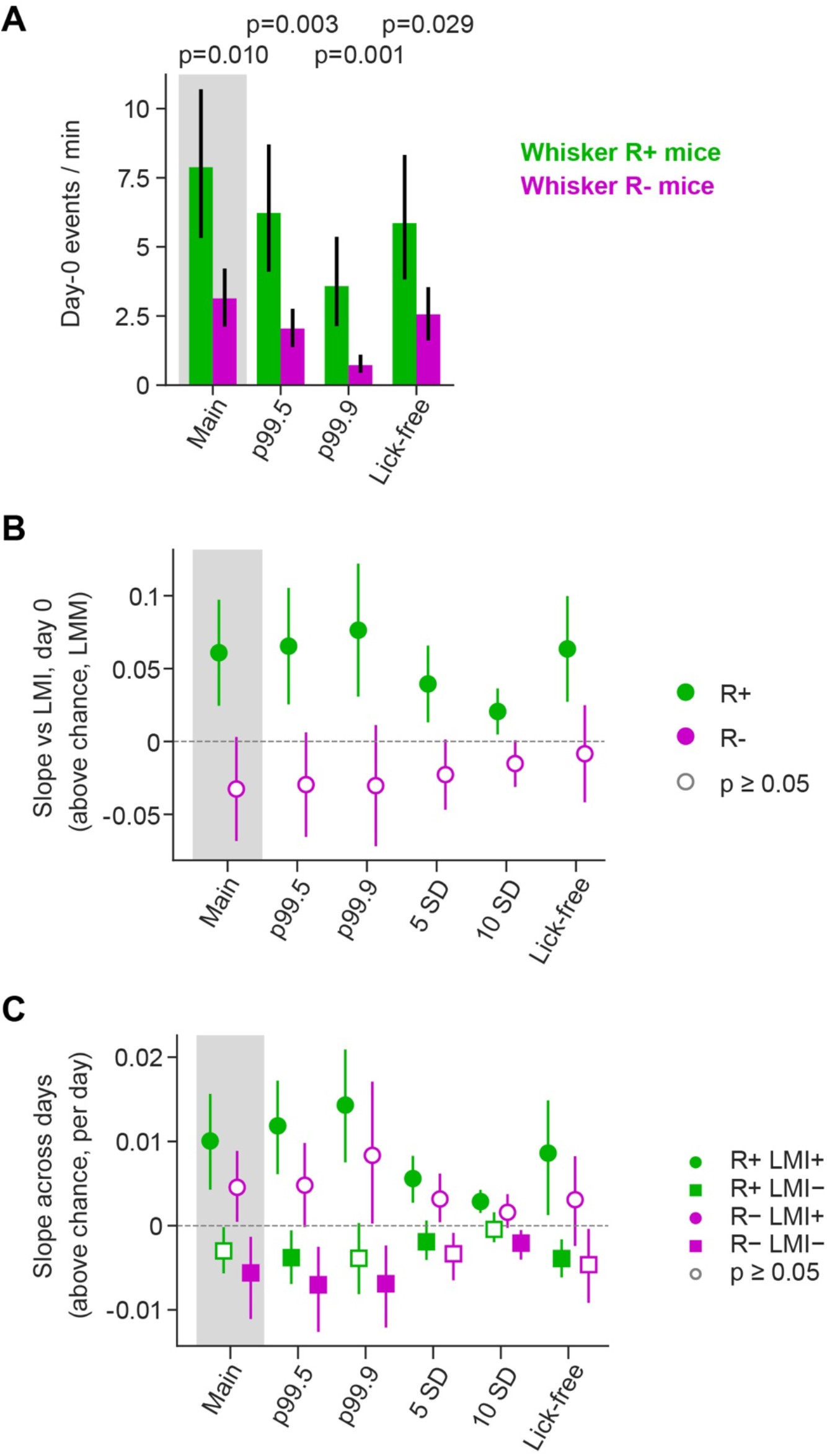
Robustness of the reactivation results to the detection threshold, the participation threshold and licking. (**A**) The analysis presented in the main figures (grey band) uses a detection threshold at the 99^th^ percentile of the surrogates, a participation threshold of 2.5 noise SD, and the whole duration of the correct-rejection trial (-1 to 6 s). To test for robustness, we changed the detection thresholds to the 99.5^th^ percentile (p99.5) and 99.9^th^ percentile (p99.9) (events re-detected), or analysed only the lick-free part of the trial (-1 to +1 s; Lick-free). Day 0 reactivation rate in R+ mice (n = 19 mice) and R-mice (n = 16 mice). Bars, mean ± 95% CI; p-values, Mann-Whitney U test. (**B**) Slope of Day 0 participation above chance vs LMI (as in Figure 6E). To test for robustness, we changed the detection thresholds to the 99.5^th^ percentile (p99.5) and 99.9^th^ percentile (p99.9) (events re-detected), participation thresholds of 5 noise SD (5 SD) and 10 noise SD (10 SD), or analysed only the lick-free part of the trial (-1 to +1 s; Lick-free). Fixed effect of the linear mixed-effects model with its 95% CI, for R+ (green) and R-mice (magenta). (**C**) Same as panel B, but for slope of participation above chance across days (as in Figure 6D) for LMI+ (circles) and LMI-neurons (squares) of R+ (green) and R-mice (magenta). Mean ± 95% CI across mice of the per-mouse slopes; Wilcoxon signed-rank test against 0. In panels B and C, filled markers, p < 0.05; open markers, p ≥ 0.05. Participation in held-out neurons, in the mice with at least three Day 0 events at the 99^th^ percentile (18 R+, 14 R-mice; 13 R-mice for LMI+ neurons).

